# Divergent cancer etiologies drive distinct B cell signatures and tertiary lymphoid structures

**DOI:** 10.1101/2020.05.29.123265

**Authors:** Ayana T Ruffin, Anthony R Cillo, Tracy Tabib, Angen Liu, Sayali Onkar, Sheryl Kunning, Caleb Lampenfeld, Irina Abecassis, Zengbiao Qi, Ryan Soose, Umamaheswar Duvvuri, Seungwon Kim, Steffi Oesterrich, Robert Lafyatis, Robert L Ferris, Dario AA Vignali, Tullia C Bruno

## Abstract

Current immunotherapy paradigms aim to reinvigorate CD8^+^ T cells, but the contribution of humoral immunity to antitumor immunity remains understudied^1,2^. Head and neck squamous cell carcinoma (HNSCC) is caused by either human papillomavirus (HPV+) or environmental carcinogens (i.e. tobacco and alcohol; HPV–)^3,4^. Here, we demonstrate that HPV+ HNSCC patients have transcriptional signatures of germinal center (GC) tumor infiltrating B cells (TIL-Bs) and spatial organization of immune cells consistent with GC-like tertiary lymphoid structures (TLS), both of which correlate with favorable outcomes in HNSCC patients. Further, our single-cell RNAseq data also indicate that GC TIL-Bs are characterized by distinct waves of gene expression consistent with dark zone, light zone and a transitional state of GC B cells. High-dimensional spectral flow cytometry permitted in depth characterization of activated, memory and GC TIL-Bs. Further, single cell RNAseq analysis and subsequent protein validation identified a role for semaphorin 4a (Sema4a) in the differentiation of GC TIL-Bs and indicated that expression of Sema4a was enhanced on GC TIL-Bs and within GC-like TLS in the TME. Thus, in contrast to some reports on the detrimental role of TIL-Bs in human tumors, our findings suggest that TIL-Bs play an instrumental role in antitumor immunity^5,6^. Novel therapeutics to enhance TIL-B responses in HNSCC should be prioritized as a compliment to current T-cell mediated immunotherapies.

## Introduction

Immunotherapy targeting the programmed cell death protein 1 (PD1) pathway is approved by the Food and Drug Administration for treatment of several metastatic or unresectable cancers including head and neck squamous cell carcinoma (HNSCC), but only ∼20% achieve a clinical benefit, highlighting the need for new therapeutic targets^1,7^. Tumor infiltrating B cells (TIL-B) represent a possible new target to compliment T cell-based immunotherapies, as they are frequent in many human tumors and positively correlate with favorable patient outcomes^8– 11^.Specifically, increased presence of TIL-B has been reported in cancers caused by environmental exposure to carcinogens (i.e., tobacco, alcohol, UV exposure) such as lung cancer and melanoma as well as cancers caused by viral infection such as hepatocellular carcinoma (HCC) and Merkel cell carcinoma (MCC)^10,12–15^. HNSCC offers a unique avenue to study TIL-Bs in the tumor microenvironment (TME) as HNSCC cancer can be caused by both exposure to environmental carcinogens or infection with high-risk human papillomavirus (HPV) ^3^. Patients with HPV+ HNSCC have historically had better outcomes compared to HPV– patients^16,17^. While the mechanisms underlying this difference in outcomes remains unknown, TIL-B are more frequent in HPV+ versus HPV– HNSCC^9,18,19^. Understanding B cell phenotypes and the spatial organization of immune populations in the TME of patients in both virally and carcinogen induce cancers will provide critical insight into ways in which TIL-Bs can be leveraged to enhance antitumor immunity.

Tertiary lymphoid structures (TLS) are immune aggregates with varying degrees of organization that form outside of secondary lymphoid organs (SLOs) in response to chronic inflammation or infection^20,21^. TLS are characterized by organization patterns similar to SLOs with defined T cell zones, B cell rich follicles and mature dendritic cells (DCs)^22,23^. TLS have been shown to also correlate with increased patient survival in many human tumors^24,25^. Recent studies have demonstrated that the presence of B cells and TLS in melanoma, renal cell carcinoma, sarcoma, and HNSCC are associated with better responses to immune checkpoint blockade (ICB)^10,11,26,27^. However, TLS are quite heterogeneous structures^28^, and the composition of TIL-Bs within these structures has not been fully elucidated. Characterization of TLS in the TME, including their composition, spatial organization, maturity, and phenotypes of immune cells involved would provide critical insight into the roles these structures play in antitumor immunity. Additionally, understanding the factors that drive formation of TLS within the TME would permit the identification of therapeutic avenues to foster an influx of TIL-Bs into the proper spatial organization.

One feature associated with mature TLS is the formation and presence of germinal centers (GCs)^29^. GCs are typically found in SLOs and are responsible for producing affinity matured and class switched B cells that effectively recognize their cognate antigen, leading to memory B cells and durable humoral immunity. In humans, GC B cells are commonly identified as CD38^+^ IgD^−^ and transcription factor Bcl6^+^. GC B cells can be further divided into centroblasts (dark zone; DZ) and centrocytes (light zone; LZ) through expression of CXCR4 and CD86. In addition, recent studies have indicated Semaphorin 4A (Sema4a) expression on human GC B cells in SLOs^30^. However, Sema4a expression on GC TIL-B has not been previously reported in human cancer. Ultimately, GCs within TLS in the TME are indicative of maximal engagement of the humoral arm of the immune system in antitumor immune responses. In support of this, GC-like TIL-Bs were found to be increased in melanoma patients who responded to ICB^10^. Understanding the features that drive TIL-Bs toward a GC phenotype and contribute to the development and maintenance of GC-like TLS in the TME would provide a path to enhancing antitumor immunity in patients.

Given the recent appreciation for TIL-Bs in the TME, we hypothesized that GC TIL-B and GC-like TLS would drive a favorable survival signature in patients with HNSCC. To address this hypothesis, we transcriptionally dissected the states of B cells in the peripheral blood (PBL) and tumors of HNSCC patients by performing scRNAseq analyses, characterized subpopulations of B cells by high-dimensional spectral flow cytometry, and assessed the spatial localization of TIL-Bs and presence of TLS in the TME using immunohistochemistry and immunofluorescence. Overall, this study demonstrates the importance of TIL-B transcriptional signatures, phenotypes and spatial patterning within the TME of patients with HNSCC, suggesting that this understudied lineage contributes to outcome and could be clinically targeted to increase antitumor immunity.

## Results

We first analyzed scRNAseq data generated from sorted CD45+ cells (i.e. all immune cells) from a total of 63 samples, including paired PBL and TIL from 18 patients with HPV– HNSCC and 9 patients with HPV+ HNSCC **(Extended Table 1, Cohort 1)**. We first developed and validated a two-step approach to robustly identify B cells and CD4^+^ T_conv_ (**Extended Figures 1 and 2;** Methods). We then bioinformatically isolated B and CD4^+^ T_conv_ and performed Louvian clustering (Methods) to reveal a total of 21 clusters (**Figure 1a**). Next, we visualized the association between sample type and transcriptional signatures by interrogating the FItSNE embedding of cells from each sample type (**Figure 1b**; Methods)^31^. Differential localization in the FItSNE revealed distinct transcriptional profiles associated with each sample type (**Figure 1b**), and association between clusters and sample types (**Figure 1c**). Based on our cell type classifications (**Extended Figure 2**), clusters 11 through 21 were B cells (**Figure 1d**), while clusters 1 through 10 were CD4^+^ T_conv_ cells (**Figure 1e**). To ascertain the role of B cells in each cluster, we filtered gene sets from the Molecular Signatures Data Base Immunologic Signatures (C7) to eight gene sets associated with canonical B cell function. This gene set enrichment analysis revealed B cell clusters associated with naïve (clusters 11, 15, 16), switched memory (clusters 12, 13, 14, 19), GC B cells (cluster 17 and 18) and plasma cells (clusters 20 and 21) (**Figure 1f**). We observed statistically significant enrichment of GC TIL-Bs and plasma cells in the TME (**Extended Figure 3a**). Interestingly, GC TIL-Bs and GC B cells from healthy tonsils were overlapping, suggesting that there is little difference between GC signatures despite being within the TME versus SLOs. We also investigated CD4^+^ T_conv_ and identified a cluster that was strongly associated with a TFH cell signature (i.e. high frequency and magnitude of *CXCR5, PDCD1, ICOS, CXCL13* expression; **Figure 1g**). These data ultimately revealed increased GC TIL-Bs in HPV+ patients and increased plasma cells in HPV– patients. Further, a TFH signature was more pronounced in HPV+ disease. To assess whether B cell signatures were clinically significant, we utilized bulk mRNAseq expression data available through The Cancer Genome Atlas (TCGA; Methods). Briefly, we scored each patient for enrichment of B cell signatures derived from our data, then determined if these gene signatures were associated with progression free survival (PFS). Overall, high B cell infiltrate, high enrichment for GC B cells, and high enrichment for memory B cells were positively associated with longer PFS (HR from 0.35 to 0.46; p from 0.003 to 0.062; **Figure 1h**). Conversely, a high frequency of plasma cells trended toward shorter PFS (HR=2.0, p=0.15; **Figure 1h**). We also found that enrichment scores for GC B cells from the light zone (LZ) were strongly correlated with those for TFH cells (rho=0.59, p<0.0001; **Extended Figure 3b**). Taken together, these data suggest that TIL-Bs in the HPV+ TME are productively activated and receive CD4^+^ T cell help (TFH).

**Figure 1:**
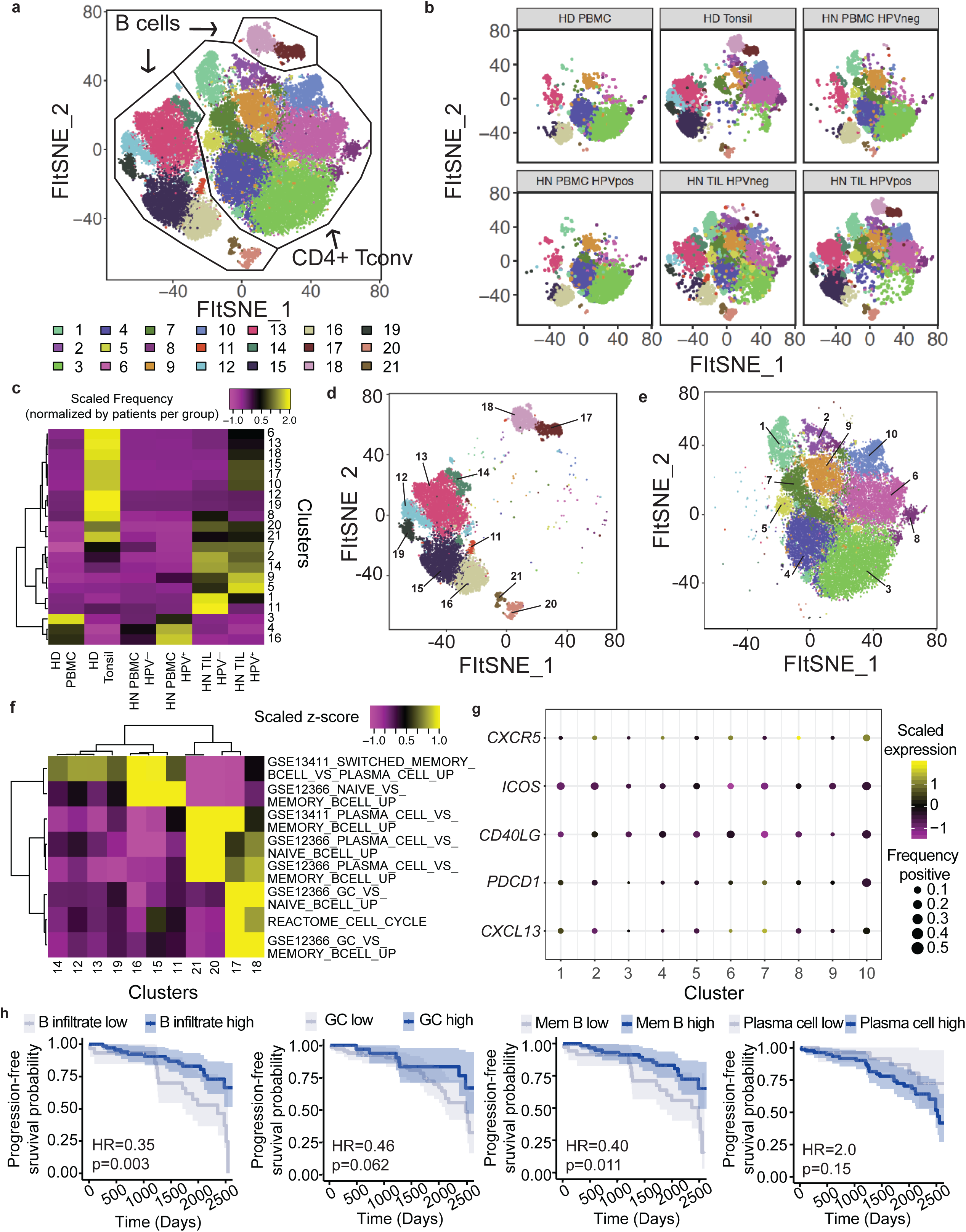
Differences in tumor infiltrating B cell and helper CD4+ T cells between HPV- and HPV+ HNSCC contribute to survival. **a**. Unsupervised clustering of 16,965 B cells and 30,092 helper CD4^+^ T cells (total of 47,057 cells) from all samples. B cells (clusters 11-21) and CD4^+^ helper T cells (1-10) form distinct groups. **b**. Same FItSNE plot as (**a**) but showing clusters by sample type. There are differences in enrichment between clusters for the different sample types, for example healthy donor tonsils and HPV+ TIL have cluster 17 and 18 which are largely absent from PBMC, while both HPV- and HPV+ TIL have B cell clusters 20 and 21. **c**. Heatmap showing the frequencies of cells recovered from each cluster by sample types, where the frequencies of cells were normalized by the number of patients assessed in each group. Tonsil samples were strong enriched for specific clusters, while HPV- and HPV+ TIL had unique patterns of cells recovered from each cluster. Statistical assessment of observed versus expected cell frequencies are detailed in **Supplementary Figure 3. d-e**. FItSNE plot (**d**) showing the clusters containing B cells from (**a**), and the associated gene sets associated with specific functions for each cluster (**e**). Canonical B cell lineages, including naïve, switched memory, plasma cells and germinal center B cells were recovered. Interestingly, cells from HPV+ patients had GC B cells, while these cells were largely absent from TIL of HPV-patients. HPV-patients had a higher frequency of naïve and memory B cells. **f-g**. FItSNE plot (**f**) showing the CD4^+^ helper T cells from (**a**), and a dot plot highlighting the present of cells with a T follicular-helper signature (cluster 10). The size of the dot is related to the frequency of cells positive for each marker, while the color is related to the magnitude of gene expression. **h**. Gene set enrichment from gene sets derived from our scRNAseq analysis were used to stratify HNSCC patients based using bulk mRNAseq expression profiles available from The Cancer Genome Atlas (TCGA), which were then used to assess progression-free survival. B cell infiltrate, germinal center B cells and memory B cells were positively associated with survival, while plasma cells were negatively associated with survival.

**Figure 2:**
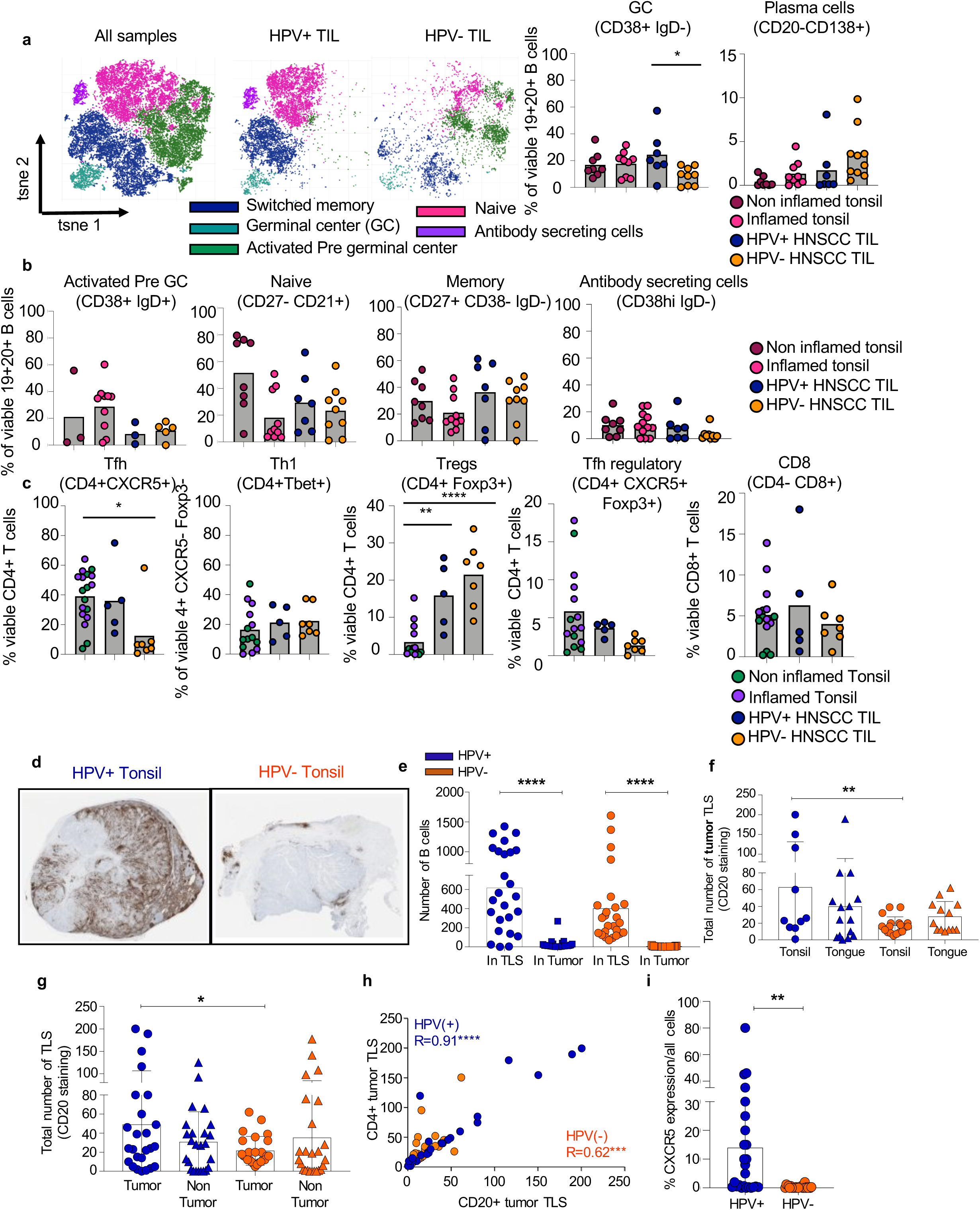
High dimensional flow cytometry and immunohistochemistry reveal distinct TIL-B phenotypes and increased tertiary lymphoid structures in HPV+ HNSCC. **a**. tSNE plots of all B cells collected HPV+ and HPV-HNSCC TIL and HNSCC PBMC analyzed using Cytofkit R program. HNSCC TIL (n=4), HNSCC PBMC (n=8). Cells are colored based on 5 populations identified using R phenograph. CD27, IgD, CD38, CXCR4, CD86, IgM surface markers were used to identify the 5 clusters. Bar plot showing frequencies of germinal center B cells and plasma cells in healthy tonsil (n=8), tonsillitis (n=9), HPV+ TIL(n=6), HPV-TIL (n=9). *P=0.02 Students 2-sided t test **b**. Bar plot showing the frequency of pre-germinal center B cells, naïve B cells, switched memory B cells and antibody secreting cells. **c**. Bar plot showing frequencies of Tfh, Tfhreg, Treg, Th1 and CD8 T cells in healthy tonsil (n=8), tonsillitis (n=10), HPV+ TIL (n= 5), HPV-TIL (n=7).*P=0.01,**P=0.004,****P<0.0001. One way ANOVA **d**. Representative CD20+ IHC on HPV (+) and HPV(-) HNSCC tumors from tonsil and tongue (4x magnification). **e**. B cells are predominantly contained within TLS compared to the tumor bed regardless of HPV status. Three areas of each patient section (n=50, 25 HPV+, 25 HPV-) were selected by the pathologist for countable B cell infiltrate in the tumor bed compared to a TLS (non GC or GC). The three areas for each patient were then averaged and subsequently graphed to reflect B cell infiltrate in the patient tumor compared to a TLS. ****P< 0.0001, Student’s 2-sided t test. **f**. Total number of tumor TLS are increased in HPV+ disease regardless of site. A HNSCC-specific pathologist identified TLS structures by organization of B cells via CD20 single-plex IHC in both patient tumor and non-tumor tissue. Enumeration of tumor TLS was parsed out by site of tumor within the oropharyngeal space (tonsil vs. tongue). Total numbers from n=50, 25 HPV+, 25 HPV(-) were graphed. **P< 0.01, Student’s 2-sided. **g**. Total number of tumor TLS are increased in HPV+ patients, however, non-tumor TLS numbers are equivalent in HPV+ and HPV- disease. Counting was done as described in (**f**). Total numbers from n=50, 25 HPV+, 25 HPV- were graphed. *P< 0.05, Student’s 2-sided t test. **h**. CD20^+^ and CD4^+^ TLS correlate in HPV+ and HPV-HNSCC patients. Total tumor TLS were independently counted for CD20^+^ and CD4^+^ by a HNSCC pathologist. Total numbers were tabulated and statistically compared (n=50, 25 HPV+, 25 HPV-). ****P< 0.0001, ***P< 0.001, non-parametric Spearman correlation. **i**. TLS-related marker, CXCR5, is increased in HPV+ HNSCC tumors. CXCR5 was scored by a HNSCC pathologist for the total percent expression of the markers across all cell types (n=50, 25 HPV+, 25 HPV-).***P< 0.001, **P< 0.001, Student’s 2 sided t test.

**Figure 3:**
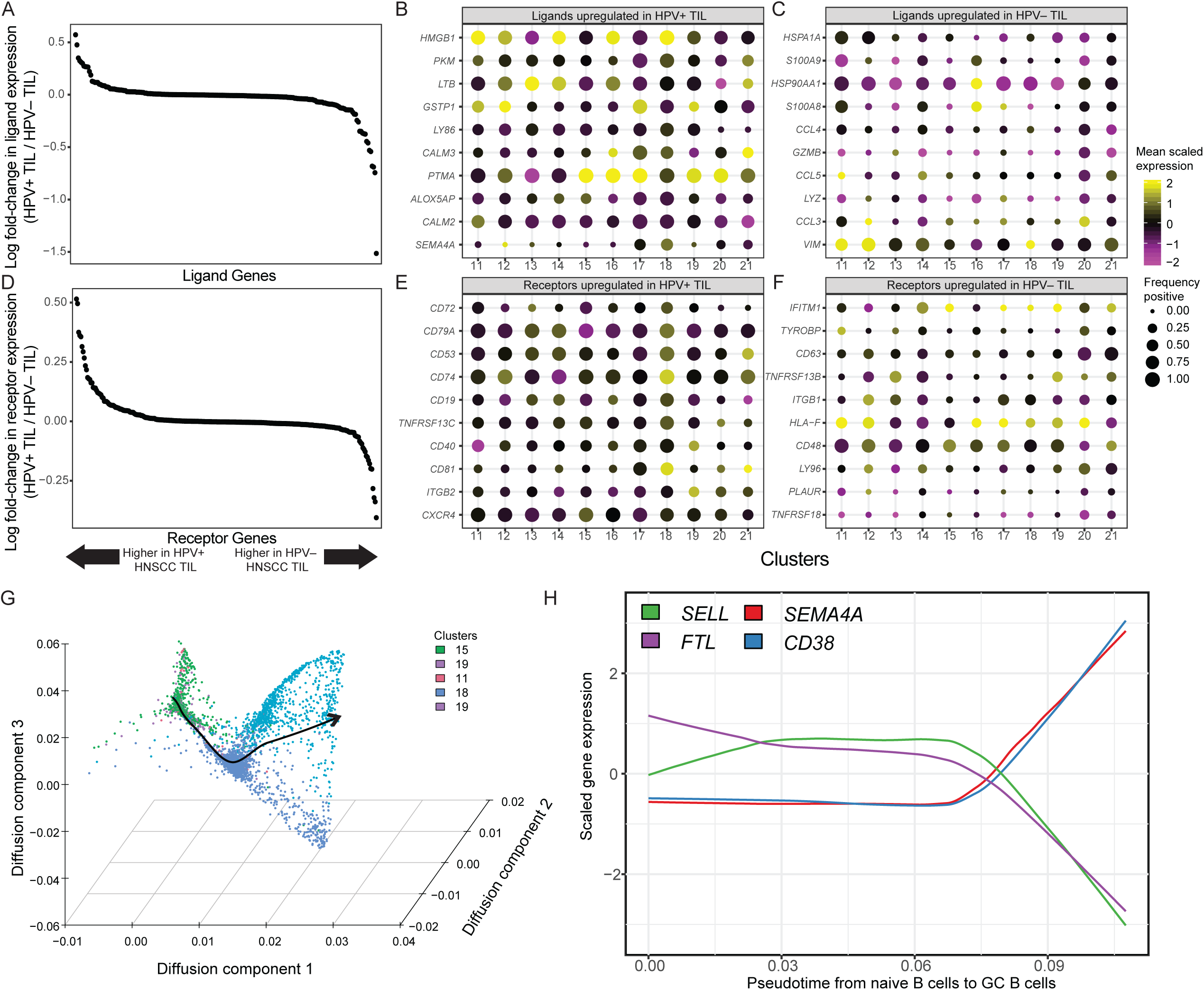
Differentially expressed ligands and receptors in HNSCC and modeling of GC differentiation identify *SEMA4A* as associated with development and maturation of GC. **a**. Differential expression of ligands by B cells in the TME of HPV– and HPV+ HNSCC. **b**. Number of cells expressing ligands and magnitude of expression in HPV+ TIL-B by cluster. Consistent with GC B cell and formation of TLS, *LTB* was one of the top expressed ligands across HPV+ TIL-B. *SEMA4A* expression was largely restricted to clusters 17 and 18, which are GC TIL-B. **c**. Expression of top ligands by HPV-TIL-B included several chemokines (*CCL4* and *CCL5*). **d**. Differential expression of receptors by B cells in the TME of HPV– and HPV+ HNSCC. **e**. Top receptors expressed by HPV+ TIL-B including genes associated with GC function including *CD40* and *CXCR4*. **f**. Top receptors in HPV– B cells included *CD63*, which is associated with downregulation of CXCR4 and is suppressed by Bcl6. **g**. Diffusion map embedding of B cell associated with a lineage spanning naïve and GC B cells identified by slingshot (Methods). B cells are shown by their clusters identified in Figure 1, and the line connecting the clusters denotes the differentiation trajectory with increasing pseudotime. **h**. Association between gene expression dynamics and differentiation from naïve to GC TIL-B shows that SEMA4A is expressed along with CD38 as naïve B cells progress to GC B cells, while SELL and FTL have the opposite expression dynamics and are downregulated during progression from naïve to GC B cells.

Given the differences in transcriptional profiles between TIL-Bs from HPV+ and HPV– HNSCC, we performed bulk B cell receptor (BCR) sequencing via Adaptive (**Extended Figure 4**; Methods). This analysis revealed no differences in measures of clonality or V-, D-, or J-gene usage between BCRs from HPV– and HPV+ HNSCC, suggesting that TIL-Bs may recognize tumor antigens in both types of HNSCC, but only receive adequate signals to support organization into GC for maximal humoral immunity in HPV+ HNSCC.

**Figure 4:**
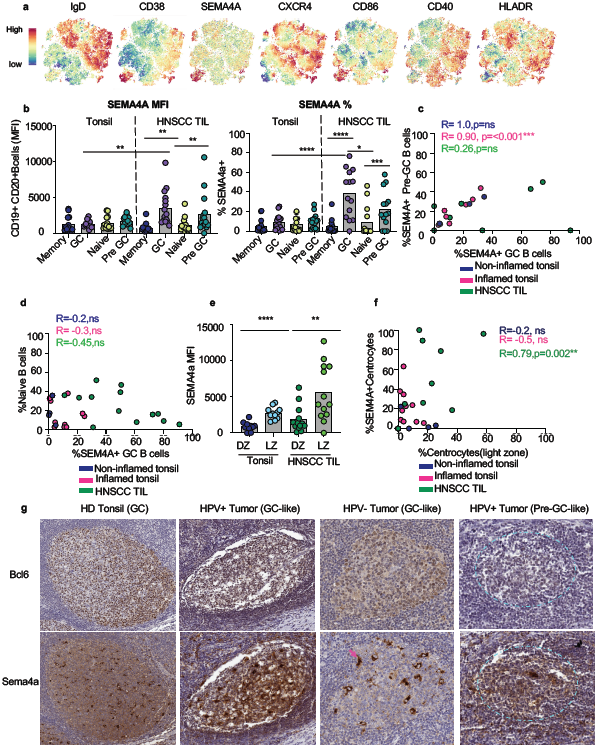
SEMA4a expression is increased in GC TIL-Bs in TLS in HNSCC. **a**. Individual feature plots demonstrating expression level of canonical markers used to identify B cell subpopulations in Figure 2a. **b**. Bar plot showing mean fluorescence intensity (MFI) of SEMA4a on B cell subsets. Bar plot showing frequencies of Sema4a positivity on B cell subsets Statistical analysis by one-way ANOVA followed by Tukeys multiple comparisons test. **P= 0.002,****P<0.0001,*P=0.04,***P=0.0003. **c**. Scatter plot comparing the frequency of SEMA4a+ Pre GC-B cells to SEMA4a GC B cells. Statistical analysis by Spearman correlation. ***P<0.001. **d**. Scatter plot comparing the frequency of SEMA4a+ GC-B cells to naive B cells. Statistical analysis by Spearman correlation. **e**. Bar plot showing MFI of SEMA4a on dark zone and light zone GC B cells. Statistical analysis by Students-T test (Mann Whitney). ****P=0.0001, **P=0.002. **f**. Scatter plot comparing the frequency of SEMA4a^+^ light zone GC-B cells to total light zone GC B cells. Scatter plot comparing the frequency of GC-B cells to T follicular helper T cells. Statistical analysis by Spearman correlation. **P=0.002. **g**. Representative IHC for Bcl6 and Sema4a in HNSCC patients. Bcl6 and Sema4a expression was compared in HPV+ and HPV-HNSCC patients to HD tonsil. Pink arrow is pathological characterization of macrophage. Blue dotted circle represents a pre-GC like TLS.

As transcriptional analysis revealed differential enrichment of TIL-Bs in HPV+ and HPV– HNSCC, we developed a spectral cytometry panel (Methods) to validate our findings at the protein level and to determine if there were any additional alterations in TIL-B subpopulations in HNSCC. We first quantified frequencies of TIL-Bs versus plasma cells in HNSCC primary tumors (Extended Table 2, Cohort 2), which revealed a significant increase in CD19^+^CD20^+^ TIL-Bs compared to plasma cells in the TME (**Extended Figure 5a-b**). Next, we utilized our spectral cytometry panel to perform unsupervised clustering of B cells on two offsets of HNSCC patients (**Cohort 2**). In the first set of patients within cohort 2, we identified five B cell clusters: naïve B cells (CD38^-^IgD^+^ CD27^-^), switched memory B cells (CD38^-^IgD^-^ CD27^+^), GC B cells (CD38^+^IgD^-^ SEMA4a^+^), activated pre-GC B cells (CD38^+^IgD^+^), and antibody secreting cells (CD38^hi^) (**Figure 2a, Extended Table 2, Cohort 2**). Consistent with the first set of patients, we identified similar B cell clusters in the second set of patients within cohort 2 but with an additional cluster of atypical memory B cell (atMBCs) (**Extended Figure 6a**). The frequency of atMBCs is highly variable in HNSCC patients, which may explain the prevalence of this population in one set of patients in cohort 2. In our first spectral cytometry analysis, TIL-Bs were predominantly associated with HPV+ HNSCC TIL, while activated pre-GC were found in HPV– HNSCC TIL (**Figure 2a**). B cells from HNSCC PBL were predominantly naïve, switched memory or activated pre-GC (**Figure 2a, Extended Figure 6a-c1**). In the second analysis, HNSCC TIL-B were mostly switched and atMBC, with some GC B cells suggesting that these patients are HPV-. (**Extended Figure 7a**). Despite the key differences we observed in this initial analysis, we wanted to quantify the frequency of these populations in additional HNSCC patients. Thus, we used traditional flow cytometry gating on cohort 2 to quantify the B cell subsets observed in the unsupervised clustering (**Extended Figure 7a**). This revealed that GC TIL-Bs were significantly increased in HPV+ HNSCC (**Figure 2b**), and that plasma cells trended towards being more frequent in HPV– HNSCC. Because our transcriptional analysis of CD4^+^ T cells in HNSCC tumors revealed an increased T follicular helper (TFH) cell signature in HPV+ HNSCC, we sought to interrogate the frequencies of CD4^+^ T_conv_ lineages (i.e. TFH, TH1, regulatory TFH, T_reg_) present in HNSCC patients. We observed a trend towards increased TFH frequencies in HPV+ HNSCC compared to HPV– tumors (**Figure 2c**), but TH1 cells were not significantly different. Regulatory TFH (CXCR5^+^ Foxp3^+^) were increased in tonsils but not significantly different between HPV+ and HPV-tumors (**Figure 2c**). T_reg_ were significantly increased in HPV– HNSCC patients compared to tonsil, and CD8^+^ T cell frequencies were similar (**Figure 2c**).

**Figure 5:**
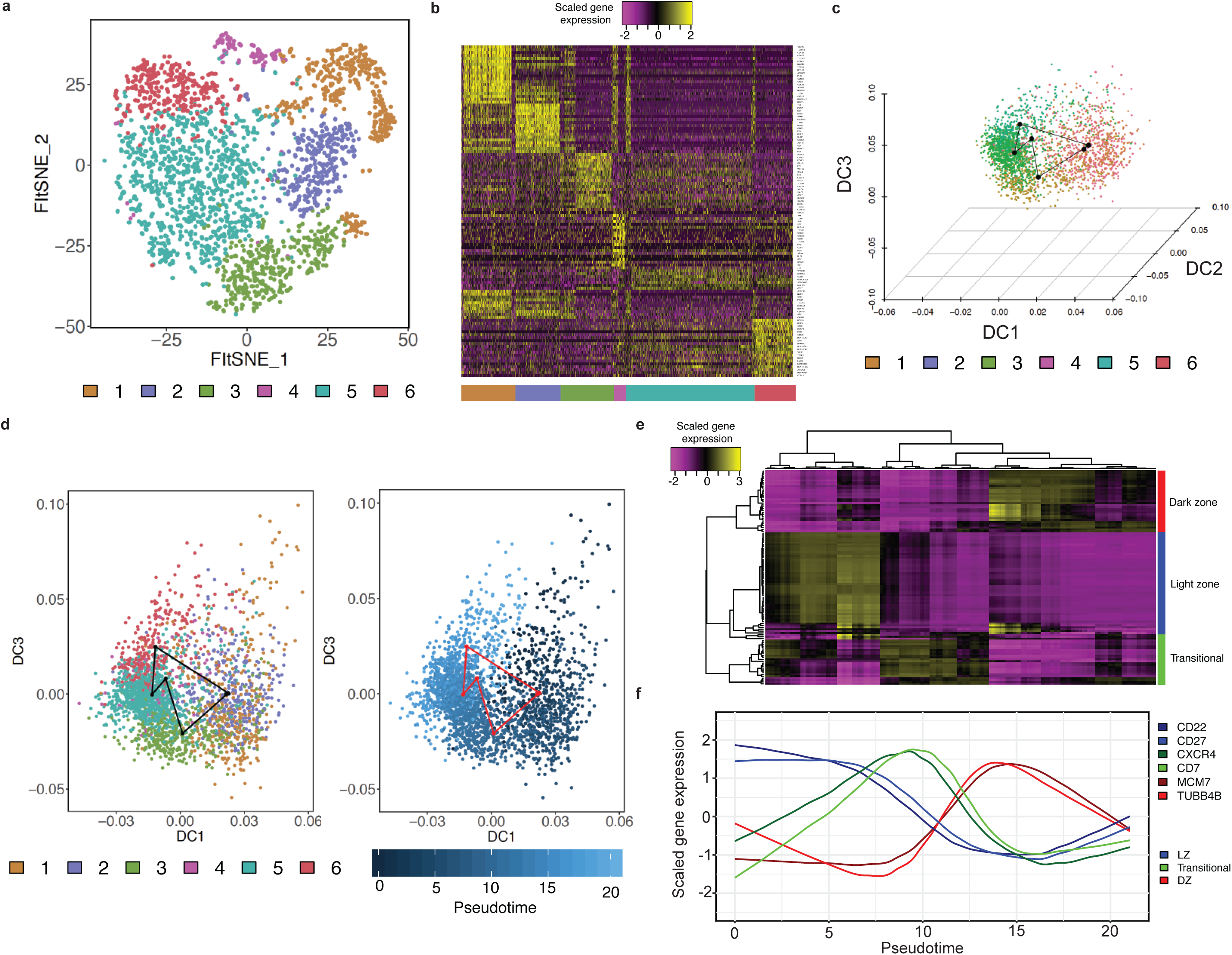
Cyclical pseudotime modeling of germinal center B cell reactions reveals waves of gene expression. **a**. FItSNE showing clusters of germinal center B cells (i.e. clusters 17 and 18 from Figure 1A/D). Louvain clustering reveald 6 clusters within the germinal center. **b**. Heatmap showing the top 20 differentially expressed genes across the 6 clusters from (**a**). **c**. Three-dimensional diffusion map embedding of germinal center B cells, which cells colored by their cluster identities from (**a**). Black dots represent the centroid of each cluster, and the lines connecting the black dots represent the circular path through germinal center reactions. **d**. DCs 1 and 3 captured most the information required to reconstruct the circular trajectory of germinal center B cells (left panel). Pseudotime order of cells from inferred by fitting the equivalent of a nonparametric principal component from the center of the trajectory using the assumption that the data is on a closed curve (right panel). This revealed a pseudotime ordering progressing through the clusters identified in (**a**). **e**. Loess regression was used to fit curves for the top 20 differentially expressed genes from (**b**) as a function of pseudotime inferred in (**d**). Genes were found to cluster into 3 distinct groups by fit with pseudotime time, suggesting distinct temporal regulation of expression in the germinal center. Loosely, these clusters of genes can be defined as dark zone, light zone and transitional genes between defined dark and light zones. **f**. Marker genes derived from (**e**), with scaled gene expression plotted as a function of time. Blue genes correspond to light zone (LZ) GC B cells, green genes correspond to B cells moving between LZ and dark zone (DZ) GC B cells, and red genes correspond to DZ GC B cells.

**Figure 6:**
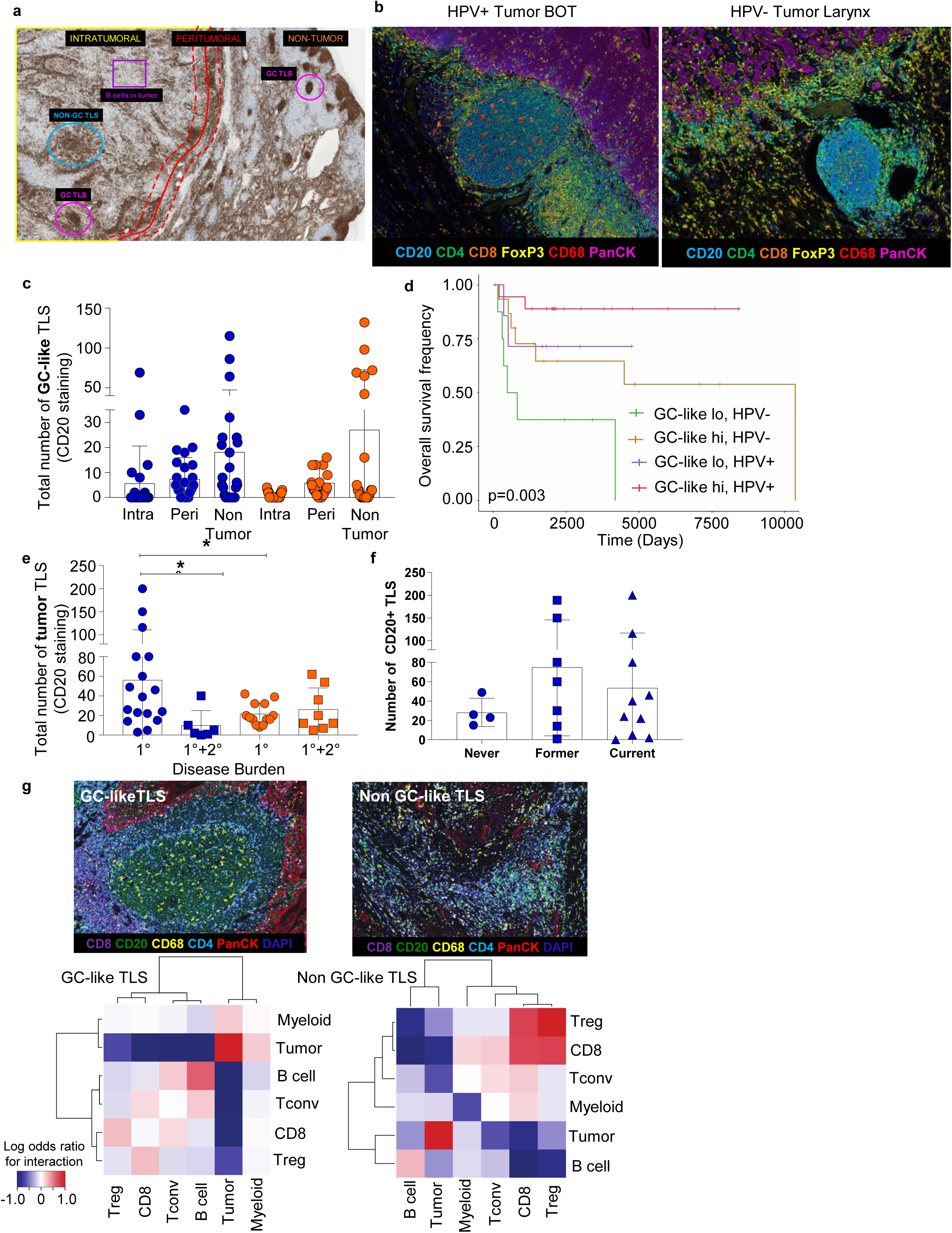
Increased GC-like TLS within HPV+ HNSCC patients correlate with increased patient survival. **a**. Annotated tumor section stained for CD20 via single-plex IHC from a HNSCC patient (20x magnification). Annotations for tumor (intratumoral and peritumoral) and non-tumor areas are indicated. Examples of non-GC-like TLS and GC-like TLS are encircled (blue and pink, respectively). An example area of the tumor bed is annotated (purple square) that would be considered TIL-B infiltration in the tumor bed which is not within a TLS (non-GC-like or GC-like). **b**. Representative Vectra staining for GC-like TLS within HPV+ and HPV-HNSCC tumors (20x magnification). Tissue sections were stained for PanCK (tumor), CD4, CD8, FoxP3 (Tregs), CD20 (B cells), and CD68 (macrophages). Seven-plex IF images were unmixed using inForm and visualized using FIJI (Methods). BOT= base of tongue **c**. GC-like TLS are increased intratumorally (intra) and peritumorally (peri) in HPV+ HNSCC patients. Tumor TLS quantification was split into intratumoral vs. peritumoral and compared again to non-tumor TLS and analysis was refined to those TLS with a GC as described in **(a)**. Differences in intra vs. peri GC-rich TLS were not statistically significant, however, they trended toward an increase in HPV+ HNSCC patients. **d**. GC-like TLS in the tumors of HPV+ and HPV-HNSCC patients correlate with increased patient survival. Cox proportional hazard was used to evaluate overall survival based on high versus low frequencies of GC-rich TLS and HPV status (p=0.003, logrank test). The hazard ratio for high versus low GC-rich TLS was 0.32, and the hazard ratio for HPV+ versus HPV-was 0.27. **e**. Total number of tumor TLS are increased in HPV+ patients that do not progress to secondary disease. Total tumor TLS (via CD20^+^ staining) were compared by patients that had only primary disease (1°) vs. primary and secondary disease (1°+2°) (as defined as recurrence at the same site). n=50, 25 HPV+, 25 HPV-.*P< 0.05, Student’s 2 sided t test. **f**. Total number of tumor TLS are increased in former and current smokers that are also HPV+. Total tumor TLS (via CD20^+^ staining) were compared in HPV+ patients that were never smokers vs. former or current smokers. **g**. Cell-cell neighborhoods in GC-like TLS are distinct compared to non-GC-like TLS. Seven-plex IF images were unmixed using inForm and visualized using FIJI (Methods). Top panels show a GC-like TLS (left) and a non-GC-like TLS (right). Bottom panels show the odds ratio of proximity to other cell types (Methods), with red representing a high probability of interaction with a given cell type and blue a low probability of interaction. The left bottom panel shows that B cells and CD4^+^ Tconv have a high probability of interacting in the GC-like TLS, while B cells were predominantly interacting with themselves and tumor cells in the non-GC-like TLS.

Although frequencies of cells quantified by flow cytometry are informative, evaluating spatial localization of cells *in situ* within the TME is an orthogonal approach that contextualizes the TME in which immune cells are located. We utilized a separate cohort (**Extended Table 3, Cohort 3**) with significant patient follow up for these locational studies. We first used single-plex immunohistochemistry (IHC) to evaluate the number and location of TIL-Bs within different areas of the oropharynx. We observed that B cells predominantly infiltrated TLS regardless of HPV status and that TLS formation was dictated by HPV status regardless of tissue sites i.e. tonsil vs. tongue (**Figure 2d-f**). Next, we evaluated frequencies of TLS in the tumor versus outside the tumor in HPV– and HPV+ HNSCC (**Figure 2g**). HPV+ tumors had a higher frequency of TLS within the tumor, and the number of CD4^+^ T cells and TIL-Bs in TLS were strongly correlated (**Figure 2h**). Finally, we found a higher frequency of CXCR5^+^ immune cells (consistent with a TFH CD4^+^ T_conv_ infiltrate) in HPV+ TIL versus HPV– TIL (**Figure 2i**), confirming that TLS likely foster GC reactions in the TME. Taken together, these flow cytometric and spatial data confirm that GC B cells and CD4^+^ TFH are present within TLS and are more frequently found in HPV+ HNSCC.

To better understand differences between TIL-B in HPV– versus HPV+ HNSCC, we next utilized our scRNAseq data to interrogate expression of ligands and receptors in the TME (**Cohort 1**). We found several ligands in the TME associated with each type of HNSCC (**Figure 3a**) and visualized the top 10 in each type of HNSCC (**Figure 3b-c**). Interestingly, we found that *SEMA4A* was a ligand that was enriched for HPV+ HNSCC and was largely restricted to GC B cell clusters (i.e. clusters 17 and 18, relative to other clusters). We performed a similar analysis with receptors, and found several receptors associated with GC B cells in HPV+ TIL (e.g. *CD40* and *CXCR4*), and others associated with plasma cells in HPV– TIL (e.g. *CD63* and *LY96*) (**Figure 3d-f**). We next used pseudotemporal modeling to better elucidate the dynamics of gene expression as cells progress from naïve B cells to GC B cells. These analyses are important not only to trace differentiation to GC B cells, but also organization of B cells into TLS, as naïve B cells must be pulled into a GC reaction to create a functional GC. Further, these analyses are supported by our scRNAseq and spectral cytometry as naïve B cells were the second B cell subset upregulated in HPV+ patients (**Figure 1c and 2a**). Briefly, pseudotemporal modeling can be used to reconstruct differentiation trajectories from scRNAseq data based on smooth changes in gene expression that take place across cells as they transition from one state to the next. We found a trajectory from naïve to GC B cells (**Figure 3g**), which allowed us to infer a pseudotime ordering of B cells for interrogation of the dynamics of gene expression from naïve to GC B cells. Interestingly, this analysis revealed that *SEMA4A* is associated with transition from naïve to GC B cells and shares similar dynamics of expression with *CD38* (**Figure 3h**). Conversely, *SELL* (gene for CD62L) and *FTL* have the opposite dynamics and are downregulated during transition from naïve to GC B cells (**Figure 3h**). Taken together, this analysis revealed that *SEMA4A* expression is enriched in GC TIL-Bs, and the temporal expression of *SEMA4A* is associated with differentiation into tissue resident, GC TIL-Bs.

With the finding that expression of *SEMA4A* on TIL-B in HNSCC patients was tightly restricted to GC B cells, we next sought to interrogate whether Sema4a has a similar expression pattern at the protein level on TIL-B (**Cohort 2**). Indeed, Sema4a was co-expressed with CD38 as in the transcriptomic data (**Figure 4a**). Further, we found that Sema4a mean fluorescence intensity (MFI) and frequency was significantly increased on GC and elevated on activated pre-GC TIL-Bs compared to GC and activated pre-GC B cells in healthy donor tonsil via our high dimensional flow cytometric data (**Figure 4a-b and Extended Figure 6b**). In addition, Sema4a MFI and frequency was significantly increased on GC TIL-Bs compared to memory or naïve TIL-Bs in HNSCC tumors (**Figure 4b**). Lastly, we observed an increase in costimulatory molecules such as CD40 and CD86 on activated pre-GC TIL-Bs compared to naïve TIL-Bs in HNSCC tumors (**Figure 4a and Extended figure 6b-d**), which we expect to be upregulated on B cell populations like GC and activated pre-GC for optimal antigen presentation. Pseudotemporal ordering in our scRNAseq data suggested that *SEMA4A* expression is increased during differentiation towards GC, meaning *SEMA4A* may play a role in the progression of activated pre-GC B cells. To interrogate this, we assessed whether there was a correlation between Sema4a+ activated pre-GC B cells and Sema4a+ GC B cells and found a trend towards positive correlation between the two groups (**Figure 4c**). We also observed an inverse correlation between Sema4a+ GC and naïve B cells in healthy tonsil and tonsillitis, and HNSCC tumors although not significant. (**Figure 4d**). Overall, these data suggest that Sema4a plays a role in development and maturation of B cells into GC B cells.

B cells entering the GC reaction begin in the dark zone (DZ) where they undergo expansion and somatic hypermutation^32,33^. Centroblasts then follow a CXCL13 gradient to enter the light zone (LZ) where they capture antigen presented on follicular dendritic cells (FDCs) which they present to T follicular helper (TFH) cells in order to undergo selection^33^. Since we observed significantly less GC TIL-Bs in HPV– HNSCC tumors, we sought to determine if there were any additional aberrations in Sema4a expression on GC B cell subsets in HNSCC tumors. Specifically, we assessed expression on DZ or LZ GC TIL-Bs. Sema4a was significantly expressed on LZ GC B cells in tonsil and HNSCC tumors (**Figure 4e**). Further, Sema4a+ LZ GC TIL-Bs positively correlate with the frequency of total LZ GC TIL-Bs (**Figure 4f**). This suggest Sema4a could be important in both the development of GC B cells and the interactions between LZ GC B cells and TFH cells in normal and tumor tissues. Using IHC, we confirmed the presence of Sema4a and co-expression of the canonical GC transcription factor Bcl6 with Sema4a in tonsils (**Figure 4g**). Interestingly, Sema4a appears to be a more robust marker of GC-like TIL-Bs in the TME of HPV+ HNSCC (**Figure 4g**). Sema4a is also more pronounced in HPV-HNSCC GC-like TLS, but is more restricted to macrophages (pink arrow) compared to TIL-Bs, whereas in HPV+ HNSCC, it is on both immune cells. Finally, we have observed precursor cells in HPV+ tissues that express Sema4a but not Bcl6, consistent with an activated pre-GC phenotype (**Figure 4f**). Taken together, these data demonstrate that Sema4a is associated with both activated pre-GC and GC B cells in tonsil and the TME of patients with HNSCC, suggesting a new role for SEMA4a in the development and maintenance of GC-like TLS in ectopic sites of inflammation.

Since a better understanding of GC reactions has implications for anti-tumor immunity and effective humoral immunity in infection and vaccination, we performed an in-depth transcriptional dissection of GC reactions. To achieve this, we first bioinformatically isolated GC B cells and re-clustered them to reveal more subtle differences within the canonical GC populations (**Figure 5a**). This analysis revealed 6 clusters with distinct gene expression patterns (**Figure 5a-b**). Typical pseudotime algorithms assume a linear differentiation trajectory, but with GC B cells we expect a cyclical process as B cells toggle between LZ and DZ interactions for optimal B cell maturity. Thus, we developed an approach to capture the cyclical nature of this process by first embedding cells in a diffusion space, yielding a cyclical topology (**Figure 5c** and Methods). We then connected each cluster via their centroids, and fit a principal curve to infer a pseudotime score for each cell in the GC (**Figure 5d**). We then evaluated genes associated with GC progression, and identified not only DZ and LZ reactions, but also a novel transitional state for TIL-Bs within our cyclical GC model (**Figure 5e**). When viewed as a function of pseudotime, we found 3 distinct waves of expression associated with each of these GC states within the cyclical process (**Figure 5f**). The first phase consisted of expression of canonical LZ genes such as *CD22* and *CD27*, followed by a wave of transitional genes consisting of *CXCR4* and *CD7*, followed by a final wave of cell cycle genes which are consistent with the proliferative nature of DZ B cells. A complete understanding of the transitional state of GC B cells will contribute to the signals that lead to egress from GC reactions, factors that contribute to the cycling between DZ and LZ, and key cues that are necessary for a bonified GC reaction in the TME.

To complement the transcriptional analysis on GC reactions in HNSCC tumors, we evaluated the number of GC-like TLS in HNSCC tumors, as GCs are paramount for maximal B cell immunity^33^. In counting GC-like vs. non-GC-like TLS in the tumor, we found elevated GC-like TLS in HPV+ and HPV-tumors (**Figure 6a-b, Extended Table 3, Cohort 3**). However, these GC-rich TLS were increased intratumorally and peritumorally in HPV+ patients (**Figure 6c**). Of note, an intratumoral increase in GC-like TLS has not been previously demonstrated in other human tumors. Further, more GC-like TLS in the tumor correlated with increased survival in both HPV+ and HPV– disease (**Figure 6d**), but more discretely in HPV+ disease, most likely due to better overall survival in these patients^16^. In addition, we revealed that HPV+ HNSCC patients with increased disease burden (i.e. primary and secondary disease) had significantly less tumor TLS in their primary disease compared to those individuals with primary disease alone (**Figure 6e**). This suggests that tumor TLS are important for reducing tumor recurrence at the same site of the primary tumor (secondary disease). We also found that former and current smokers with the HPV+ cohort of patients had increased TLS compared to never smokers (**Figure 6f**). This indicates the importance of another environmental cues in TLS formation in cancer. Finally, we analyzed the key cell-cell neighborhoods in GC-like vs. non-GC-like TLS in HNSCC (**Figure 6g**). In GC-like TLS, TIL-Bs interact with other TIL-Bs and CD4^+^ T_conv_ TIL, which is in line with the working definition of an active GC. Interestingly, an evaluation of a non-GC like TLS in HNSCC revealed that TIL-Bs were not frequently next to CD4^+^ T_conv_, and instead CD8^+^ TIL and T_regs_ were implicated as a dominant interaction. These results demonstrate that in GC-like TLS, the spatial patterning becomes distinct from well-infiltrated tumors where immune cells are found in aggregates.

## Discussion

In this study, we sought to perform an in-depth analysis of B cells in the TME of patients with HNSCC, with the goal of improving our understanding of the immunobiology of B cells and the potential role they have in generating baseline antitumor immune responses. Our study integrated new technical approaches across three cohorts of patient samples (n=124) and suggests that not only higher numbers of TIL-Bs, but also the specific phenotype and localization of TIL-Bs in the TME contribute to overall survival. Specifically, we are the first to report that Sema4a^+^ GC TIL-Bs and GC-like TLS are increased in HPV+ HNSCC patients compared to HPV-.Further, we also identified CD4^+^ TFH in the TME of HNSCC, which complements findings in breast and colorectal cancer^34–36^. The correlation we observed between LZ B Cells and TFH in the TME extends this finding further, demonstrating the importance of crosstalk between CD4^+^ T cells and GC TIL-Bs and the need for CD4^+^ T cell help for GC TIL-B survival in the TME of HNSCC. Our single-cell transcriptional characterization of TIL-B populations uncovered numerous states of B cells in the TME and revealed distinct differences between HPV+ and HPV-HNSCC. These differences should be considered in the development of a B cell-focused immunotherapy for HNSCC.

B cells are a heterogenous population with phenotypically and functionally distinct subsets. Thus, characterization of TIL-B phenotypes in treatment naïve patients is a critical first step in the development of B cell-focused immunotherapies. However, B cell targeted therapies may need to enhance certain subsets of B cells while inhibiting others, necessitating more dissection of the change in TIL-B phenotypes following therapy. For example, in melanoma, patients who did not respond to standard of care immunotherapy i.e. anti-PD1 and/or anti-CTLA4 had significantly more naïve B cells than responders^10^. In this case, would depleting naïve B cells increase patient response or would driving naïve B cells to differentiate and enter GC reactions be effective? Our data would suggest that this is a viable therapeutic consideration as naïve TIL-Bs, GC TIL-Bs, CD4^+^ TFH and GC-like TLS were all significantly increased in HPV+ HNSCC patients. However, a functional assessment of TIL-B subpopulations is needed to better inform potential targeting strategies. There are a multitude of ways in which B cells can contribute to antitumor immunity, and it will be important to link B cell subsets with specific antitumor function.

One function for TIL-Bs that is definitively correlated with increased survival and immunotherapeutic response in cancer patients is their role in TLS^15,22,23,26,37^. TLS formation and maintenance in tumors is an active area of investigation. Early studies reveal that common mechanisms of lymphoid organogenesis such as the presence of inflammatory cytokines and interactions of immune cells with tissue-resident stromal cells such as fibroblasts and mesenchymal cells are important for TLS initiation ^22–24,38,39^. Our study identifies a potential mechanism for TLS formation in tumors through the identification of SEMA4a expression on GC TIL-Bs within TLS. *SEMA4A* is a membrane bound glycoprotein that is important for T cell co-stimulation and an important driver of Th2 responses in humans, and was recently found to be expressed on human GC B cells in SLOs^30^. Further, SEMA4a can interact with non-immune receptor Plexin D1 which is expressed on endothelial cells and immune receptor T cell, Ig domain, mucin domain-2 (Tim-2) and neuropilin-1 (NRP1) expressed by T cells^40–44^. Thus, SEMA4A may play a central role in generating immune aggregates via TIL-B interactions with endothelial and T cells. In fact, CD4^+^ TFH express high levels of NRP1^43^, which is the main CD4^+^ T cell subset where we observe correlations with GC TIL-Bs. Future studies should more thoroughly characterize the factors that lead to the creation of effective TLS, or conversely the factors that inhibit TLS formation in the TME, especially because TLS are both predictive of ^37,45,46^ and correlated with response to immunotherapy^10,11,26^.

Current immunotherapeutic regimens aim to re-invigorate exhausted CD8^+^ TIL within the TME ^47^. Overall, our findings suggest that engagement of humoral immunity in treatment naive patients is associated with better outcomes. Focusing on amplifying early activation of TIL-Bs into GC-like TLS is a potentially paradigm-shifting step towards new immunotherapies. For example, we found that Sema4a may be a better marker of both early-stage and functional TLS in the TME compared with the canonical GC B cell marker Bcl6. As such, determining ways to drive Sema4a expression on TIL-B and determining which ligands are required to nucleate TLS is an obvious next step for B cell mediated immunotherapy development. These findings are likely to stimulate future studies involving Sema4a in other cancers that have reported GC-TIL-Bs such as lung cancer and melanoma^10,15,29^.In addition, formation of GC-like TLS both peritumorally and intratumorally is paramount for increased patient survival and are increased in virally induced HNSCC. Thus, our study has implications for other virally induced cancers such as HCC, MCC, and cervical cancer where the presence of GC-TIL-Bs has not yet been reported. Future studies should seek to evaluate how viral infection impacts the development and maintenance of GC-TIL-B and GC-like TLS in virally induced cancers. Further, additional environmental factors (i.e. the microbiome of the oral cavity and oropharynx) should be queried in future studies. Lastly, improved analysis of spatial relationships will be paramount as our data suggest that GC biology within TLS is associated with favorable anti-tumor immunity. Beyond cancer, our dissection of B cell biology can inform strategies aimed at enhancing vaccine responses, or conversely disrupting the generation of B-cell mediated immune activation to suppress autoimmunity. Ultimately, this study highlights the significance of phenotypes and spatial patterns of TIL-Bs in both virally and carcinogen induced cancer and suggests that therapeutic enhancement of antitumor humoral immunity should be paired with current immunotherapeutic platforms.

## Author contributions

TCB conceived the project. DAAV and TCB obtained funding. ATR, ARC, DAAV, and TCB interpreted data and wrote the manuscript. ATR performed flow cytometry experiments and analyzed flow cytometry data. ARC performed scRNAseq experiments, analyzed scRNAseq data and immunofluorescence data, and performed statistical analyses. SK and IA performed flow cytometric experiments. SaO performed immunofluorescence staining (within the lab of co-mentor StO). CL performed analysis and quantification of IHC images. AL performed IHC image analysis and interpretation (with TCB). RLF, UD, SK and RJS identified patients and collected specimens. RLF provided feedback and clinical interpretation of the data. TT and ZQ performed single-cell RNAseq experiments; RL provided input on single-cell RNAseq experimental design and library preparation. All authors reviewed and approved the manuscript.

## Author information

The authors declare no competing financial interests.

## Materials and methods

### Patient cohorts

Several patient cohorts were used for various aspects of this manuscript. Figures 1, 3, 5 and Extended Figures 1-4 used the patient cohort described in Extended Table 1. This cohort has been previously described in detail^48^ Figure 2 and 4 and Extended Figures 5-7 used the patient cohorts described in Extended Tables 2. Figure 6 used the patient cohort described in Extended Table 3. All patients provided informed written consent prior to donating samples for this study, and the study was approved by the Institutional Review Board (University of Pittsburgh Cancer Institute, Protocol 99-069).

### Blood and tissue processing

Peripheral blood was obtained by venipuncture into and collected into tubes containing EDTA coagulant. Blood was processed into PBMC by Ficoll-Hypaque density gradient centrifugation. Briefly, whole blood was diluted and layered over Ficoll-Hypaque, followed by centrifugation at 400xg for 20 minutes with the brake set to off. PBMC were then collected and washed in complete RPMI (i.e. RPMI 10% fetal bovine serum and 1% penicillin/streptomycin).

Tissues were collected from either HNSCC patients undergoing resection as treatment or sleep apnea or tonsillitis patient undergoing tonsillectomy. Tissues were collected directly into collection media (i.e. complete RPMI + 1% amphotericin B) in the operating room and were processed as soon as possible following surgery. For transcriptional analysis, samples were processed within 2 hours of collection. Sample processing consistent of manually dissociating tumor tissue into approximately 1 mm pieces, then washing with cRPMI and passing the suspension over a 100 uM filter. The filter was then washed with cRPMI, and the cells were centrifuged at 500xg for 5 minutes. If significant numbers of red blood cells were present, red blood cell lysis was performed as per the manufacturer’s instructions (BD Pharm Lyse).

### Flow cytometry-based cell sorting

For experiments requiring cell sorting, cells were first stained in PBS with 2% FBS and 1 mM EDTA for 15 minutes, followed by centrifugation at 500xg for 5 minutes and staining with viability factor in PBS for 15 minutes. Cells were then centrifuged again, resuspended in PBS with 2% FBS and 1 mM EDTA, and sorted using a MoFlo Astrios High Speed Sorter (Beckman Coulter). Sort cells were collected directly in cRPMI. For single-cell RNAseq analysis, live CD45+ cells were sorted by using Fixable Viability Dye eFluor780 (eBioscience) and CD45 conjugated to PE (Biolegend, clone HI30).

### Single-cell RNAseq Library Preparation and Sequencing

Immediately following sorting, cells were centrifuged for 5 minutes at 500xg and were resuspended in PBS with 0.04% BSA. Cells were then counted using the Cellometer Auto2000 (Nexcelom), and loaded into the 10X Controller (10X Genomics) as per the manufacturer’s instructions. Following bead/cell emulsification, cDNA synthesis was performed as per the manufacturer’s instructions (10X Genomics). cDNA was then purified by SPRI-bead selection as per the manufacturer’s instructions, and cDNA was then amplified and fragmented for library generated followed by 12 cycles of PCR amplification. The library quality was determined by Bioanalyzer analysis and concentration by KAPA qPCR DNA Quantification. Libraries were then pooled and sequenced on a NextSeq500 (University of Pittsburgh Genomics Research Core) using a high-output kit, targeting a read depth of 100,000 reads/cell.

### Processing and clustering of single-cell RNAseq data

Following sequencing, raw Illumina reads were demultiplexed based on i7 indices (10X Genomics) using the mkfastq command of the CellRanger suite of tools (10X Genomics). Demultiplexed FASTQs were then aligned to the human genome (GrCH38) using the count command of CellRanger to generate cell/barcode matrices. Cell/barcode matrices were then read into Seurat (v2.3.4) for downstream analysis.

Clustering was performed as an initial analysis step for several scRNAseq datasets using the workflow popularized by Seurat. Briefly, raw reads were normalized for library size per cell and log-transformed. Highly variable genes were identified and selected, followed by scaling and center of data as well as regression out technical variables (i.e. number of genes per cell, percent of reads aligning to ribosomal genes per cell and percent of reads aligning to mitochondrial reads per cell). These scaled and centered expression values were then used as input into a principal component analysis to reduce the dimensionality of the data. The top principal components that explained the most variance in the dataset were heuristically selected as input for the fast interpolation-based t-SNE^31^ and the Louvian-based clustering algorithm implemented in Seurat.

### Identification of cell types in single-cell RNAseq

We initially sorted and sequenced all cells of the hematopoietic lineage (i.e. CD45+ cells), and were therefore needed to robustly identify B cells and CD4^+^ T_conv_ for downstream in-depth analysis. We did this using a two-step semi-supervised identification strategy. This strategy consisted of first identifying core transcriptional programs of the major lineages of the immune compartment. To do this, we downloaded publicly available single-cell RNAseq data of sorted immune lineages (10X Genomics; https://www.10xgenomics.com/resources/datasets/). We then clustered these cell populations as described above to identify lineage-specific clusters. Once these clusters were identified, we performed differential gene expression analysis using a Wilcoxon rank sum test to identify the top 20 genes associated with each cluster. These genes were defined as the core transcriptional profile of each lineage. We then used these genes as gene sets to test individual cells for enrichment of each immune lineage. Briefly, we used the log-fold change in gene expression as a metric and input these fold-changes into the Wilcoxon rank sum test for genes in each core lineage set versus genes outside that set, deriving a gene set score and p-value for each gene set for each cell. The core lineage gene set associated with the lowest p-value for each cell was then applied as that cell type. Following this test for each cell, we then examined clusters of cells in aggregate, and identified each cluster by the most common cell type enriched within that cluster. We then compared this two-step method (i.e. single-cell gene set enrichment testing and identification followed by aggregate identification of clusters) to the ground truth for each of the clusters know to be a sorted cell lineage using a confusion table from the R package caret.

### Single-cell RNAseq analysis of B and CD4^+^ T_conv_

To identify B cells and CD4^+^ T_conv_ from our dataset of all hematopoietic cells, we applied the two-step method described above. This yielded pure populations of B and CD4^+^ T_conv_, which were then clustered as described above. We next evaluated the enrichment of cells from a given sample type in each cluster by dividing the frequency of observed cells over expected cells in each cluster. The expected frequency of cells was calculated by assuming cells from each sample group were evenly divided across the clusters. Analysis of variance was used to determine if the cell enrichment across groups was statistically significant. Gene set enrichment analysis was performed using a variance inflated Wilcoxon rank sum test^48^ for B cells, using input gene sets available from the Molecular Signatures Database (C7 Immunology Gene Sets). These gene sets were then pre-filtered for those relevant to B cells, and then curated based on specific B cell signatures. Differential gene expression analysis using a variance inflated Wilcoxon rank sum test (described above) was used to identify gene expression patterns across clusters.

### Survival analysis using The Cancer Genome Atlas

To determine if our gene sets were relevant for survival, we utilized bulk RNAseq data for HNSCC patients available through the TCGA and create an enrichment score for each signature from each patient as previously described^48^. Briefly, we used the top 200 differentially expressed genes from specific clusters of B cells to determine an enrichment score for genes in that gene set versus genes outside that gene set using a Kolmogorov-Smirnov test. We then stratified patients based on high versus low enrichment scores and performed Cox proportional hazards regression (see statistical analysis below).

### Pseudotime analysis of B cells

Clustering analysis is useful for grouping cell types based on similar gene expression patterns but does not capture information related to developmental trajectories of cells. To assess developmental trajectories, we first embedded cells in a low-dimensional diffusion map (e.g. performed non-linear dimensionality reduction^49^. We then used the R package slingshot^50^ to infer a pseudotime for each cell along the developmental trajectory, and to infer individual trajectories. To evaluate whether genes were statistically associated with pseudotime, we performed LOESS regression using the R package gam, where we fit gene expression as a function of pseudotime along each trajectory. We focused on the trajectory that was characterized by progression from naïve B cells to germinal center B cells.

For pseudotime analysis of germinal center B cells, slingshot could not be used since it assumes a linear trajectory. Germinal center B cells are in a cycle between light and dark zones, and therefore require pseudotime inference based on a cyclical process. Therefore, a principal curve was fit along the circular trajectory to infer the pseudotime of each cell in this process. Genes were once again investigated for their relationship to pseudotime and were clustered based correlation of gene expression over pseudotime.

### Adaptive B cell receptor Sequencing

Adaptive Biotechnologies’ immunoSEQ platform was used to perform a survey of B cell receptors (BCRs) from HNSCC patients. Total DNA was isolated from cryopreserved snap frozen tumor tissues using the QIAGEN DNeasy Blood and Tissue Kit and was used as input for the immunoSEQ platform. Analysis was performed using Adaptive’s analysis interface.

### Surface and intracellular antibody staining of patient and healthy donor cells

Single cell suspensions from either HNSCC tissue, tonsillar tissue, HNSCC PBL or healthy donor PBL were stained with fluorescently labeled antibodies for 25 mins at 4°C in PBS (Thermo Fisher) supplemented with 2% FBS (Atlanta Biologicals) and 0.01% azide (Thermo Fisher) (FACS buffer). Antibodies purchased from Biolegend or BD against: CD19 (HIB19), CD20 (2H7), CD21 (Bu32), CD27 (0323), IgM (G20-127), IgD (JAG-2), CD138 (M-115), LAIR1 (NKTA255), FcRL4 (413D12), FcRL5 (509f6), CD40 (5C3), CD86 (IT2.2), CXCR4 (12G5), CD38 (HB-7), CD11c (3.9), CD70 (Ki-24), CD39(A1), CD85j (GHI/75), CD95 (DX2), CXCR3 (1C6), CXCR5 (JS52D4), Tbet (4B10), CD178 (NOK-1), CD73 (AD2) were used to stain Cohort 2 in **Figure 2-a-b**. Additionally, CD45 (H130), CD19 (HIB19), CD20 (2H7), CD21 (Bu32), CD27 (0323), HLADR (307619), CD40 (5C3), CD86 (IT2.2), CD69 (FN50), CD70 (Ki-24), CXCR4 (12G5), SEMA4A (5E3), CD22 (HIB22), LAIR1 (NKTA255), VISTA (MIH65), FcRL4 (413D12), FcRL5 (509f6), PDL1 (29E.2A3), IgM (G20-127), IgD (JAG-2), CD138 (M-115),CD23(M-L233)and CD38 (HB-7) were used to stain Cohort 2 in **Extended Figure 6 and some patients in Figure 2-a-b**. To exclude myeloid, T and NK cells from B cell analysis, the following antibodies labeled with the same fluorophore were used: CD14 (63D3), CD11c (3.9), CD11b (ICRF44), CD66b (G10F5), TCR α/β (IP26) and CD56 (5.1H1). Cohort 2 in **Extended figure 5** were stained with antibodies against CD19 (SJ25C1), CD20 (2H7), CD27 (O323), CD21 (HB5), CD38 (HB-7), CD86 (IT2.2), CD40 (5C3), CD138 (M115), PD1 (eBioJ105), PDL1 (29E.2A3), LAG-3 (3DS223H), IgA (IS11-8E10), CD69 (FN50), and HLADR (G46-6). Cells were also stained with a T cell-specific panel using the following antibodies against CD4 (RPA-T4), CD8 (RPA-T8), CXCR5 (JS52D4), Tbet (4B10), FoxP3 (PCH101), Bcl6 (K112-91), CD45RA (H1100), NRP1 ((12C2), CD27(O323), ICOS (C398.4A), CCR7 (G043H7), CD25 (BC96), (**Figure 2c**). Cells were stained using Fixable Viability Dye (eBioscience) in PBS to exclude dead cells. For intracellular transcription factor staining cells were fixed using fixation/permeabilization buffer (eBioscience) for 20 mins at 4°C the washed with permeabilization buffer (eBioscience). Cells were then stained with fluorescently labeled antibodies. Flow cytometry measurements were performed on an LSR-II flow cytometer (BD) or Cytek Aurora (Cytek). All data were analyzed using FlowJo.

### High dimensional spectral cytometry

Data were analyzed using Cytofkit, a mass cytometry package for R programing software as previously described^51^. Briefly, B cells were pre-gated on CD19+CD20+ in flow jo. Surface markers of interest were selected, and FCS files were exported from Flowjo and imported in to the Cytofkit package. Expression values for each selected surface marker are extracted from each FCS file and transformed using automatic logicle transformation (autoLgcl). The software combines each expression matrix using a selected method. For the data in this manuscript we selected, *ceil* which takes a specified number of cells to include in the analysis without replacement from each FCS file. Next, we selected the Phenograph clustering algorithm. R phenograph identified 28 clusters. Cluster identification was then determined by assessing surface marker expression in each cluster using t-SNE visualization. We used the expression of CD27, CD38, IgD, IgM, CXCR4, and CD86 to reduce the clustering to five biologically meaningful clusters.

### Single-plex immunohistochemistry

Fresh tissues were formalin-fixed immediately followed surgical resection and were then embedded in paraffin. Tissues were processed as previously described^48^. Briefly, fixed tissues were then slide mounted, de-paraffinized using xylene and ethanol, and then re-fixed in formalin for 15 minutes followed by antigen retrieval. Slides were stained with the following antibodies: CD20 (Clone L26, Invitrogen), CD4 (Clone SP35, ThermoFisher), CXCR5 (Clone D6L36, Cell signaling), Tbet (Clone 4B10, Abcam). Quantification of cells and TLS were performed by a HNSCC pathologist. Specifics of these quantifications are outlined in Figure Legends and definitions of a TLS were consistent across three independent pathologists.

### Immunofluorescence analysis

Fresh tissues were formalin-fixed immediately followed surgical resection and were then embedded in paraffin. Tissues were processed as previously described^48^. Briefly, fixed tissues were then slide mounted, de-paraffinized using xylene and ethanol, and then re-fixed in formalin for 15 minutes followed by antigen retrieval as per the manufacturer’s instructions (Perkin Elmer). Blocking was performed for 10 minutes, followed by incubation with primary antibodies for 30 minutes. Secondary antibodies conjugated to horseradish peroxidases were then added for 10 minutes. Cells were stained with the following conjugated opal dyes: CD4/Opal540, CD8/Opal570, CD20/Opal520, CD68/Opal650, FOXP3/Opal620 and Pan-cytokeratin/Opal690. Cells were also counterstained with DAPI and sealed with Diamond Anti-fade mounting (ThermoFisher).

Following staining, slides were imaged as whole slide scans on the Vectra (Perkin Elmer). Regions of interest were selected from the whole slide scans, and slides were re-imaged to captures these regions at 10x magnification. Images were unmixed after scanning using inForm and Phenochart. Custom macros were written to utilize FIJI for high-throughput standardized image analysis^48^. Briefly, cells were segmented via watershed analysis in each individual channel, and cells were assigned an x- and y-position on each slide associated with their cell type. We then performed Delaunay triangulation to determine to odds of a cell interaction with another given cell type based on proximity^48,52^.

### Statistical analysis

Analysis of variance (ANOVA) followed by pairwise t-tests was used to compare more than two groups of continuous variables. Two groups of continuous variables were compared by t-tests or Wilcoxon rank sum tests were indicated. Tukey’s multiple comparisons test was performed following ANOVA where indicated. Survival analysis was performed by using Cox proportional hazards regression analysis, using either nominal values or stratifying continuous variables into nominal values. Stratification of continuous variables was performed using the “cutp” function of the R package survMisc. Correlations were performed using either Pearson’s correlation or Spearman’s correlation, as indicated. Correction for multiple comparisons using the false discovery rate was performed where appropriate. P values and false discovery rates were considered statistically significant when the two-sided type I error was 5% or less.

## Acknowledgements

The authors thank the head and neck cancer clinical team (Amy Cuda, Merida Serrano, Tina Harrison, and Denise Knoll) for collection of patient samples; the Hillman Cancer Center Flow Cytometry Core (Bratislav Janjic, Ernest Meyer, Paul Dascani) for cell sorting and help with Cytek panel design; the Health Science Core Research Facilities Genomics Research Core for Illumina sequencing; the Tissue and Research Pathology Facilities at UPMC Shadyside (Anthony Green); the University of Colorado Human Immune Monitoring Shared Research Facility, the Immunologic Monitoring and Cellular Products Laboratory at Hillman Cancer Center, and the National Surgical

Adjuvant Breast and Bowel Project (Marion Joy) for immunofluorescence imaging; and the University of Pittsburgh Center for Research Computing for computational resources. This study was funded by support through the University of Pittsburgh Cancer Immunology Training Program (T32 CA082084 to AR and ARC), the Hillman Postdoctoral Fellowship for Innovative Cancer Research (to ARC), the National Institutes of Health (P50 HNSCC SPORE CA907190 to DAAV and RLF, CDA and DRP to TCB), Eden Hall Pilot Funds (to TCB), and the NCI Comprehensive Cancer Center Support CORE grant (CA047904 to DAAV and RLF).

## Supplementary Materials

**Extended Table 1: Clinical characteristics of prospective patient cohort for single-cell RNAseq and immunofluorescence (Cohort 1)**

**Extended Table 2: Clinical characteristics of prospective patient cohort for spectral flow cytometry and protein validation (Cohort 2)**

**Extended Table 3: Clinical characteristics of retrospective patient cohort for IHC and TLS analysis (Cohort 3)**

**Extended Figure 1: Validation of the combination Wilcoxon rank sum test and clustering based method for identification of cell types**.

**Extended Figure 2: Identification of cell types from patients and controls using the combination Wilcoxon rank sum test and clustering based approach**.

**Extended Figure 3: Statistical assessment of observed versus expected number of cells in each cluster by sample types**.

**Extended Figure 4: Adaptive BCR sequencing reveals no difference in clonality or other metrics between HPV^−^ and HPV^+^ TIL**.

**Extended Figure 5: B cells are significantly increased compared to plasma cells in HNSCC patients**.

**Extended Figure 6: Additional high dimensional analysis of HNSCC cohort 2**.

**Extended Figure 7: Flow cytometry gating strategy for B cell and T cell profiling**.

## Bibliography

1. Ferris, R. L. et al. Nivolumab for Recurrent Squamous-Cell Carcinoma of the Head and Neck. N. Engl. J. Med. 375, 1856–1867 (2016).

2. Economopoulou, P., Kotsantis, I. & Psyrri, A. The promise of immunotherapy in head and neck squamous cell carcinoma: combinatorial immunotherapy approaches. ESMO Open 1, e000122 (2016).

3. Dok, R. & Nuyts, S. HPV positive head and neck cancers: molecular pathogenesis and evolving treatment strategies. Cancers (Basel) 8, (2016).

4. Spence, T., Bruce, J., Yip, K. W. & Liu, F.-F. HPV associated head and neck cancer. Cancers (Basel) 8, (2016).

5. Tang, A. et al. B cells promote tumor progression in a mouse model of HPV-mediated cervical cancer. Int. J. Cancer 139, 1358–1371 (2016).

6. Inoue, S., Leitner, W. W., Golding, B. & Scott, D. Inhibitory effects of B cells on antitumor immunity. Cancer Res. 66, 7741–7747 (2006).

7. Wu, X. et al. Application of PD-1 Blockade in Cancer Immunotherapy. Comput Struct Biotechnol J 17, 661–674 (2019).

8. Garaud, S. et al. Tumor infiltrating B-cells signal functional humoral immune responses in breast cancer. JCI Insight 5, (2019).

9. Wood, O. et al. Gene expression analysis of TIL rich HPV-driven head and neck tumors reveals a distinct B-cell signature when compared to HPV independent tumors. Oncotarget 7, 56781–56797 (2016).

10. Helmink, B. A. et al. B cells and tertiary lymphoid structures promote immunotherapy response. Nature 577, 549–555 (2020).

11. Petitprez, F. et al. B cells are associated with survival and immunotherapy response in sarcoma. Nature 577, 556–560 (2020).

12. Miller, N. J. et al. Merkel cell polyomavirus-specific immune responses in patients with Merkel cell carcinoma receiving anti-PD-1 therapy. J. Immunother. Cancer 6, 131 (2018).

13. Zur Hausen, A., Rennspiess, D., Winnepenninckx, V., Speel, E.-J. & Kurz, A. K. Early B-cell differentiation in Merkel cell carcinomas: clues to cellular ancestry. Cancer Res. 73, 4982–4987 (2013).

14. Xiao, X. et al. PD-1hi Identifies a Novel Regulatory B-cell Population in Human Hepatoma That Promotes Disease Progression. Cancer Discov. 6, 546–559 (2016).

15. Germain, C. et al. Presence of B cells in tertiary lymphoid structures is associated with a protective immunity in patients with lung cancer. Am. J. Respir. Crit. Care Med. 189, 832–844 (2014).

16. Fakhry, C. et al. Improved survival of patients with human papillomavirus-positive head and neck squamous cell carcinoma in a prospective clinical trial. J. Natl. Cancer Inst. 100, 261–269 (2008).

17. Weinberger, P. M. et al. Molecular classification identifies a subset of human papillomavirus--associated oropharyngeal cancers with favorable prognosis. J. Clin. Oncol. 24, 736–747 (2006).

18. Lechner, A. et al. Tumor-associated B cells and humoral immune response in head and neck squamous cell carcinoma. Oncoimmunology 8, 1535293 (2019).

19. Hladíková, K. et al. Tumor-infiltrating B cells affect the progression of oropharyngeal squamous cell carcinoma via cell-to-cell interactions with CD8+ T cells. J. Immunother. Cancer 7, 261 (2019).

20. Pitzalis, C., Jones, G. W., Bombardieri, M. & Jones, S. A. Ectopic lymphoid-like structures in infection, cancer and autoimmunity. Nat. Rev. Immunol. 14, 447–462 (2014).

21. Neyt, K., Perros, F., GeurtsvanKessel, C. H., Hammad, H. & Lambrecht, B. N. Tertiary lymphoid organs in infection and autoimmunity. Trends Immunol. 33, 297–305 (2012).

22. Sautès-Fridman, C. et al. Tertiary lymphoid structures in cancers: prognostic value, regulation, and manipulation for therapeutic intervention. Front. Immunol. 7, 407 (2016).

23. Sautès-Fridman, C., Petitprez, F., Calderaro, J. & Fridman, W. H. Tertiary lymphoid structures in the era of cancer immunotherapy. Nat. Rev. Cancer 19, 307–325 (2019).

24. Germain, C., Gnjatic, S. & Dieu-Nosjean, M.-C. Tertiary Lymphoid Structure-Associated B Cells are Key Players in Anti-Tumor Immunity. Front. Immunol. 6, 67 (2015).

25. Castino, G. F. et al. Spatial distribution of B cells predicts prognosis in human pancreatic adenocarcinoma. Oncoimmunology 5, e1085147 (2016).

26. Cabrita, R. et al. Tertiary lymphoid structures improve immunotherapy and survival in melanoma. Nature 577, 561–565 (2020).

27. Kim, S. S. et al. B-cells improve overall survival in HPV-associated squamous cell carcinomas and are activated by radiation and PD-1 blockade. Clin. Cancer Res. (2020). doi: 10.1158/1078-0432.CCR-19-3211

28. Kroeger, D. R., Milne, K. & Nelson, B. H. Tumor-Infiltrating Plasma Cells Are Associated with Tertiary Lymphoid Structures, Cytolytic T-Cell Responses, and Superior Prognosis in Ovarian Cancer. Clin. Cancer Res. 22, 3005–3015 (2016).

29. Siliņa, K. et al. Germinal centers determine the prognostic relevance of tertiary lymphoid structures and are impaired by corticosteroids in lung squamous cell carcinoma. Cancer Res. 78, 1308–1320 (2018).

30. Lu, N. et al. Human Semaphorin-4A drives Th2 responses by binding to receptor ILT-4. Nat. Commun. 9, 742 (2018).

31. Linderman, G. C., Rachh, M., Hoskins, J. G., Steinerberger, S. & Kluger, Y. Fast interpolation-based t-SNE for improved visualization of single-cell RNA-seq data. Nat. Methods 16, 243–245 (2019).

32. Bannard, O. et al. Germinal center centroblasts transition to a centrocyte phenotype according to a timed program and depend on the dark zone for effective selection. Immunity 39, 912–924 (2013).

33. Mesin, L., Ersching, J. & Victora, G. D. Germinal center B cell dynamics. Immunity 45, 471–482 (2016).

34. Gu-Trantien, C. et al. CD4^+^ follicular helper T cell infiltration predicts breast cancer survival. J. Clin. Invest. 123, 2873–2892 (2013).

35. Gu-Trantien, C. et al. CXCL13-producing TFH cells link immune suppression and adaptive memory in human breast cancer. JCI Insight 2, (2017).

36. Bindea, G. et al. Spatiotemporal dynamics of intratumoral immune cells reveal the immune landscape in human cancer. Immunity 39, 782–795 (2013).

37. Figenschau, S. L., Fismen, S., Fenton, K. A., Fenton, C. & Mortensen, E. S. Tertiary lymphoid structures are associated with higher tumor grade in primary operable breast cancer patients. BMC Cancer 15, 101 (2015).

38. Luther, S. A. et al. Differing activities of homeostatic chemokines CCL19, CCL21, and CXCL12 in lymphocyte and dendritic cell recruitment and lymphoid neogenesis. J. Immunol. 169, 424–433 (2002).

39. Cupovic, J. et al. Central Nervous System Stromal Cells Control Local CD8(+) T Cell Responses during Virus-Induced Neuroinflammation. Immunity 44, 622–633 (2016).

40. van der Zwaag, B. et al. PLEXIN-D1, a novel plexin family member, is expressed in vascular endothelium and the central nervous system during mouse embryogenesis. Dev. Dyn. 225, 336–343 (2002).

41. Delgoffe, G. M. et al. Stability and function of regulatory T cells is maintained by a neuropilin-1-semaphorin-4a axis. Nature 501, 252–256 (2013).

42. Renand, A. et al. Neuropilin-1 expression characterizes T follicular helper (Tfh) cells activated during B cell differentiation in human secondary lymphoid organs. PLoS One 8, e85589 (2013).

43. Battaglia, A. et al. Neuropilin-1 expression identifies a subset of regulatory T cells in human lymph nodes that is modulated by preoperative chemoradiation therapy in cervical cancer. Immunology 123, 129–138 (2008).

44. Sarris, M., Andersen, K. G., Randow, F., Mayr, L. & Betz, A. G. Neuropilin-1 expression on regulatory T cells enhances their interactions with dendritic cells during antigen recognition. Immunity 28, 402–413 (2008).

45. Lee, H. J. et al. Tertiary lymphoid structures: prognostic significance and relationship with tumour-infiltrating lymphocytes in triple-negative breast cancer. J. Clin. Pathol. 69, 422–430 (2016).

46. Dieu-Nosjean, M.-C. et al. Long-term survival for patients with non-small-cell lung cancer with intratumoral lymphoid structures. J. Clin. Oncol. 26, 4410–4417 (2008).

47. Im, S. J. et al. Defining CD8+ T cells that provide the proliferative burst after PD-1 therapy. Nature 537, 417–421 (2016).

48. Cillo, A. R. et al. Immune Landscape of Viral- and Carcinogen-Driven Head and Neck Cancer. Immunity 52, 183–199.e9 (2020).

49. Haghverdi, L., Büttner, M., Wolf, F. A., Buettner, F. & Theis, F. J. Diffusion pseudotime robustly reconstructs lineage branching. Nat. Methods 13, 845–848 (2016).

50. Street, K. et al. Slingshot: cell lineage and pseudotime inference for single-cell transcriptomics. BMC Genomics 19, 477 (2018).

51. Chen, H. et al. Cytofkit: A bioconductor package for an integrated mass cytometry data analysis pipeline. PLoS Comput. Biol. 12, e1005112 (2016).

52. Goltsev, Y. et al. Deep Profiling of Mouse Splenic Architecture with CODEX Multiplexed Imaging. Cell 174, 968–981.e15 (2018).

