## Supplemental data for "Divergent cancer etiologies drive distinct B cell signatures and tertiary lymphoid structures"

| Patient Identifier | Tumor p16 status | Tumor location | Pathologic Stage | Pathological Node | Sex | Race | Age | Tobacco Use | Alcohol Use |
|---|---|---|---|---|---|---|---|---|---|
| **Healthy donor 1** | NA | NA | NA | NA | Female | Caucasian | 34 | No | Occassional |
| **Healthy donor 2** | NA | NA | NA | NA | Male | Caucasian | 29 | No | Yes |
| **Healthy donor 3** | NA | NA | NA | NA | Female | Caucasian | 60 | Former | No |
| **Healthy donor 4** | NA | NA | NA | NA | Male | Caucasian | 63 | Former | Occassional |
| **Healthy donor 5** | NA | NA | NA | NA | Female | Caucasian | 55 | No | Occassional |
| **Healthy donor 6** | NA | NA | NA | NA | Male | Caucasian | 56 | No | Yes |
| **Healthy tonsil 1** | NA | NA | NA | NA | Male | Caucasian | 53 | No | Unknown |
| **Healthy tonsil 2** | NA | NA | NA | NA | Male | Caucasian | 34 | Former | Unknown |
| **Healthy tonsil 3** | NA | NA | NA | NA | Male | Caucasian | 28 | Yes | No |
| **Healthy tonsil 4** | NA | NA | NA | NA | Male | Caucasian | 38 | Former | No |
| **Healthy tonsil 5** | NA | NA | NA | NA | Male | Caucasian | 47 | Former | Occassional |
| **HNSCC 1** | p16- | Oral cavity | T1 | N2b | Male | Caucasian | 45 | Yes | Yes |
| **HNSCC 2** | p16- | Floor of mouth | T4 | N0 | Male | Caucasian | 50 | Yes | Yes |
| **HNSCC 3** | p16- | Floor of mouth | T4 | N0 | Female | Caucasian | 66 | Former | Unknown |
| **HNSCC 4** | p16- | Buccal mucosa | T4 | N1 | Male | Caucasian | 48 | Yes | Yes |
| **HNSCC 5** | p16- | Tongue | T3 | N2b | Male | Caucasian | 45 | Yes | Occassional |
| **HNSCC 6** | p16- | Larynx | T4 | N2c | Male | Caucasian | 43 | Yes | No |
| **HNSCC 7** | p16- | Lower gum | T4 | N3b | Male | Caucasian | 74 | Yes | No |
| **HNSCC 8** | p16- | Tongue | T3 | N2a | Female | Caucasian | 60 | No | No |
| **HNSCC 9** | p16- | Oral cavity | TX | NX | Male | Caucasian | 80 | No | No |
| **HNSCC 10** | p16- | Tongue | T3 | N2B | Male | Caucasian | 56 | Yes | Yes |
| **HNSCC 11** | p16- | Tongue | T3 | N3B | Female | Caucasian | 57 | Yes | No |
| **HNSCC 12** | p16- | Tongue | T3 | N0 | Male | Caucasian | 35 | No | No |
| **HNSCC 13** | p16- | Supraglottis | T3 | N0 | Male | Caucasian | 62 | Yes | Yes |
| **HNSCC 14** | p16- | Mandable | T4A | N2C | Female | Caucasian | 77 | Yes | No |
| **HNSCC 15** | p16- | Tongue | T3 | N0 | Female | Caucasian | 75 | Yes | Yes |
| **HNSCC 16** | p16- | Tongue | T3 | N0 | Male | Caucasian | 55 | No | Yes |
| **HNSCC 17** | p16- | Buccal mucosa | T2 | N0 | Male | Caucasian | 80 | No | No |
| **HNSCC 18** | p16- | Floor of mouth | T2 | N1 | Female | Caucasian | 69 | Yes | Yes |
| **HNSCC 19** | p16+ | Base of tongue | T2 | N2b | Male | Caucasian | 68 | No | Occassional |
| **HNSCC 20** | p16+ | Base of tongue | T1 | N2b | Male | Caucasian | 62 | Former | Occassional |
| **HNSCC 21** | p16+ | Base of tongue | T1 | N2a | Male | Caucasian | 49 | Former | Unknown |
| **HNSCC 22** | p16+ | Tonsil | T2 | N1 | Male | Caucasian | 66 | No | Occasional |
| **HNSCC 23** | p16+ | Tongue | T4 | N1 | Male | Caucasian | 61 | Yes | Yes |
| **HNSCC 24** | p16+ | Larymx | Unknown | Unknown | Male | Unknown | 15 | Unknown | Unknown |
| **HNSCC 25** | p16+ | Base of tongue | T1 | N1 | Male | Caucasian | 52 | Former | No |
| **HNSCC 26** | p16+ | Tonsil | T2 | N0 | Male | Unknown | 75 | No | No |
| **HNSCC 27** | p16+ | Tonsil | T1 | N1 | Male | Caucasian | 59 | Yes | No |

**Extended Table 1: Clinical characteristics of prospective patient cohort for single-cell RNAseq and immunofluorescence (Cohort 1)**

| Patient Identifier | Tumor p16 status | Tumor location | Pathologic Stage | Pathological Node | Sex | Race | Age | Tobacco Use | Alcohol Use |
|---|---|---|---|---|---|---|---|---|---|
| **HNSCC 1** | p16+ | Tonsil | unknown | unknown | Female | Caucasian | 55 | Yes | occasional |
| **HNSCC 2** | not evaluated | Tongue | T3 | N3B | Female | Caucasian | 57 | Yes | Yes |
| **HNSCC 3** | not evaluated | Gum | TX | N1B | Female | Caucasian | 49 | Yes | Yes |
| **HNSCC 4** | p16- | Larynx | unknown | unknown | Female | Caucasian | 52 | Yes | yes |
| **HNSCC 5** | p16- | Tongue | T3 | N2B | Female | Caucasian | 75 | Yes | yes |
| **HNSCC 6** | p16- | Tongue | T3 | N0 | Male | Caucasian | 55 | No | yes |
| **HNSCC 7** | not evaluated | Tongue | T3 | N1 | Female | Caucasian | 61 | Yes | occasional |
| **HNSCC 8** | not evaluated | Tongue | T1 | N0 | Male | Caucasian | 66 | No | occasional |
| **HNSCC 9** | not evaluated | Thyroid gland | T2 | N1B | Male | Caucasian | 53 | yes | occasional |
| **HNSCC 10** | p16+ | Base of Tongue | T0 | N1 | Male | Caucasian | 51 | yes | yes |
| **HNSCC 11** | not evaluated | Larynx | T4A | N0 | Male | Unknown | 81 | yes | No |
| **HNSCC 12** | not evaluated | Thyroid gland | T3 | N0 | Male | Caucasian | 59 | yes | yes |
| **HNSCC 13** | p16+ | Base of Tongue | T1 | N1 | Male | Caucasian | 64 | No | yes |
| **HNSCC 14** | p16- | Tongue | T3 | N3B | Male | Caucasian | 56 | yes | yes |
| **HNSCC 15** | not evaluated | Larynx- glottis | T4A | N3B | Male | Caucasian | 64 | no | yes |
| **HNSCC 16** | not evaluated | Base of Tongue | T3 | N1 | Female | Caucasian | 60 | yes | yes |
| **HNSCC 17** | p16+ | Base of Tongue | T1 | N1 | Male | Caucasian | 52 | yes | yes |
| **HNSCC 18** | p16+ | Base of Tongue | T2 | N0 | Male | Caucasian | 78 | yes | yes |
| **HNSCC 19** | not evaluated | tongue | T4A | N3B | Female | Caucasian | 60 | no | occasional |
| **HNSCC 20** | not evaluated | Mouth- anterior floo | T1 | N0 | Male | Caucasian | 58 | yes | no |
| **HNSCC 21** | not evaluated | Larynx- glottis | unknown | unknown | Female | Caucasian | 66 | yes | yes |
| **HNSCC 22** | not evaluated | Tongue | T3 | N0 | Female | Caucasian | 32 | yes | yes |
| **HNSCC 23** | not evaluated | Larynx- supraglottis | T3 | N2C | Male | Caucasian | 55 | yes | yes |
| **HNSCC 24** | not evaluated | Larynx- supraglottis | unknown | unknown | Male | Caucasian | 56 | yes | yes |
| **HNSCC 25** | p16+ | Tonsil | T1 | N2B | male | Caucasian | 36 | no | yes |
| **HNSCC 26** | not evaluated | Base of Tongue | T3 | N0 | Female | Caucasian | 49 | yes | no |
| **HNSCC 27** | p16- | mouth | T4A | N3 | Male | Caucasian | 57 | yes | no |
| **HNSCC 28** | p16- | Gum | T4A | N3B | Male | Caucasian | 58 | yes | yes |
| **HNSCC 29** | not evaluated | Tongue | T3 | N1 | Male | Caucasian | 64 | yes | occasional |
| **HNSCC 30** | not evaluated | Larynx- glottis | T3 | N0 | Male | Caucasian | 48 | yes | yes |
| **HNSCC 31** | p16+ | Base of Tongue | T2 | N1 | Male | Caucasian | 71 | no | occasional |
| **HNSCC 32** | not evaluated | Larynx | TX | NX | Male | Caucasian | 62 | yes | yes |
| **HNSCC 33** | not evaluated | mouth | T4 | N0 | Male | Caucasian | 63 | yes | yes |
| **HNSCC 34** | not evaluated | hypopharynx | TX | NX | Male | Caucasian | 78 | yes | yes |
| **HNSCC 35** | p16+ | base of Tongue | T3 | N1 | Male | Caucasian | 55 | no | no |
| **HNSCC 36** | p16+ | base of Tongue | T1 | N1 | Female | Caucasian | 55 | no | yes |
| **HNSCC 37** | p16+ | Tonsil | T1 | N2B | Female | Caucasian | 46 | yes | yes |
| **HNSCC 38** | p16+ | Tonsil | T1 | N1 | Male | Caucasian | 55 | yes | occasional |
| **HNSCC 39** | p16+ | Tonsil | T2 | N1 | Male | frican America | 57 | yes | yes |
| **HNSCC 40** | p16- | hypopharynx | T3 | N2C | Male | Caucasian | 62 | yes | yes |
| **HNSCC 41** | p16- | base of Tongue | T4A | N2C | Male | Caucasian | 47 | no | no |
| **HNSCC 42** | p16- | Larynx- supraglottis | T3 | N2C | Male | Caucasian | 61 | yes | yes |
| **HNSCC 43** | p16- | Larynx- supraglottis | T3 | N2C | Female | Caucasian | 66 | yes | occasional |
| **HNSCC 44** | p16- | Tongue | T4A | N0 | Male | Caucasian | 74 | yes | yes |
| **HNSCC 45** | p16- | Tonsil | T1 | N3 | Male | Caucasian | 56 | yes | yes |
| **HNSCC 46** | p16- | Larynx- supraglottis | T3 | N2C | Female | Caucasian | 66 | yes | occasional |
| **HNSCC 47** | not evaluated | gum | T2 | N2C | Female | Caucasian | 50 | yes | occasional |

**Extended Table 2: Clinical characteristics of prospective patient cohort for spectral flow cytometry and protein validation (Cohort 2)**

| Patient Identifier | Tumor p16 status | Tumor location | Pathologic Stage | Pathological Node | Sex | Race | Age | Tobacco Use | Alcohol Use |
|---|---|---|---|---|---|---|---|---|---|
| **HNSCC 1** | p16+ | Tongue-base | T1 | N2A | Male | Caucasian | 66 | No | Yes |
| **HNSCC 2** | p16+ | Tonsil | T1 | N2B | Male | Caucasian | 53 | Yes | Yes |
| **HNSCC 3** | p16+ | Tonsil | T1 | N1 | Male | Caucasian | 58 | No | Occasional |
| **HNSCC 4** | p16+ | Tongue | T2 | N2C | Male | Caucasian | 64 | Yes | Yes |
| **HNSCC 5** | p16+ | Tonsil | T2 | N0 | Male | Caucasian | 46 | No | Occasional |
| **HNSCC 6** | p16+ | Tongue-base | T3 | N1 | Male | Caucasian | 75 | Unknown | Unknown |
| **HNSCC 7** | p16+ | Tonsil | T2 | N2B | Male | Caucasian | 63 | Former | Yes |
| **HNSCC 8** | p16+ | Tonsil | T3 | N2B | Male | Caucasian | 69 | Former | Yes |
| **HNSCC 9** | p16+ | Tongue-base | T1 | N1 | Male | Caucasian | 60 | Unknown | Unknown |
| **HNSCC 10** | p16+ | Tonsil | T1 | N1 | Male | Caucasian | 37 | Yes | No |
| **HNSCC 11** | p16+ | Tongue-base | T2 | N0 | Male | Caucasian | 50 | Yes | No |
| **HNSCC 12** | p16+ | Tongue-base | T2 | N2 | Male | Caucasian | 60 | Unknown | Unknown |
| **HNSCC 13** | p16+ | Tongue-base | Unknown | Unknown | Male | Caucasian | 49 | Former | Yes |
| **HNSCC 14** | p16+ | Tongue | T1 | N0 | Male | Caucasian | 55 | Yes | Yes |
| **HNSCC 15** | p16+ | Tongue-base | T2 | N1 | Male | Caucasian | 61 | Yes | No |
| **HNSCC 16** | p16+ | Tonsil | T2 | N0 | Male | Caucasian | 47 | Yes | Yes |
| **HNSCC 17** | p16+ | Tongue-base | T4 | N2C | Male | Caucasian | 58 | Unknown | Unknown |
| **HNSCC 18** | p16+ | Tongue-base | Unknown | Unknown | Male | Caucasian | 50 | Yes | No |
| **HNSCC 19** | p16+ | Tonsil | T1 | N2B | Male | Caucasian | 49 | Former | Yes |
| **HNSCC 20** | p16+ | Tonsil | T1 | Unknown | Male | Caucasian | 52 | Former | Yes |
| **HNSCC 21** | p16+ | Tongue-base | T2 | N2B | Male | Caucasian | 66 | Yes | Unknown |
| **HNSCC 22** | p16+ | Tongue-base | T4 | N2B | Male | Caucasian | 38 | Yes | Yes |
| **HNSCC 23** | p16+ | Tongue-base | T2 | N1 | Male | Caucasian | 61 | Former | Former |
| **HNSCC 24** | p16+ | Tongue-base | T3 | N2B | Male | Caucasian | 57 | Former | Yes |
| **HNSCC 25** | p16+ | Tonsil | T2 | N0 | Female | Caucasian | 60 | No | No |
| **HNSCC 26** | p16- | Tongue | T3 | N2B | Male | Caucasian | 53 | Yes | Yes |
| **HNSCC 27** | p16- | Tongue-base | T3 | N2C | Male | Caucasian | 61 | Yes | Unknown |
| **HNSCC 28** | p16- | Tonsil | T1 | N0 | Female | Caucasian | 54 | Yes | Unknown |
| **HNSCC 29** | p16- | Tonsil | T2 | N2B | Male | Caucasian | 66 | Former | Yes |
| **HNSCC 30** | p16- | Tonsil | T2 | N2B | Male | Caucasian | 48 | Former | Yes |
| **HNSCC 31** | p16- | Tonsil | T2 | N2B | Male | Caucasian | 60 | Former | Yes |
| **HNSCC 32** | p16- | Tonsil | T1 | N1 | Male | Caucasian | 82 | No | No |
| **HNSCC 33** | p16- | Tonsil | T1 | N0 | Female | Caucasian | 65 | Former | Yes |
| **HNSCC 34** | p16- | Tongue | Unknown | Unknown | Male | Caucasian | 60 | No | Yes |
| **HNSCC 35** | p16- | Tongue-base | Unknown | Unknown | Male | Caucasian | 62 | Unknown | Unknown |
| **HNSCC 36** | p16- | Tongue-base | T3 | N1 | Male | Caucasian | 57 | Yes | Yes |
| **HNSCC 37** | p16- | Tonsil | T1 | N0 | Male | AA | 63 | Yes | No |
| **HNSCC 38** | p16- | Tonsil | T2 | N0 | Female | Caucasian | 60 | Former | Former |
| **HNSCC 39** | p16- | Tongue-base | T2 | N2B | Male | Caucasian | 67 | Yes | Yes |
| **HNSCC 40** | p16- | Tonsil | T4 | N2B | Male | Caucasian | 48 | Former | Yes |
| **HNSCC 41** | p16- | Tongue-base | T2 | N2 | Male | Caucasian | 70 | Yes | Yes |
| **HNSCC 42** | p16- | Tonsil | T2 | N0 | Male | Caucasian | 42 | Yes | Yes |
| **HNSCC 43** | p16- | Tonsil | T4 | N1 | Male | AA | 57 | Yes | Yes |
| **HNSCC 44** | p16- | Tongue-base | T3 | N1 | Male | Caucasian | 60 | Unknown | Unknown |
| **HNSCC 45** | p16- | Tonsil | T2 | N0 | Male | Caucasian | 65 | Yes | Yes |
| **HNSCC 46** | p16- | Tongue-base | T1 | N2 | Female | Caucasian | 54 | Former | Yes |
| **HNSCC 47** | p16- | Tonsil | T2 | N1 | Female | Caucasian | 61 | Yes | No |
| **HNSCC 48** | p16- | Tonsil | T1 | N0 | Male | Caucasian | 68 | Yes | Yes |
| **HNSCC 49** | p16- | Tonsil | T3 | N2B | Male | AA | 51 | Unknown | Unknown |
| **HNSCC 50** | p16- | Tongue-base | T1 | N1 | Male | Caucasian | 46 | Yes | Former |

**Extended Table 3: Clinical characteristics of retrospective patient cohort for IHC and TLS analysis (Cohort 3)**

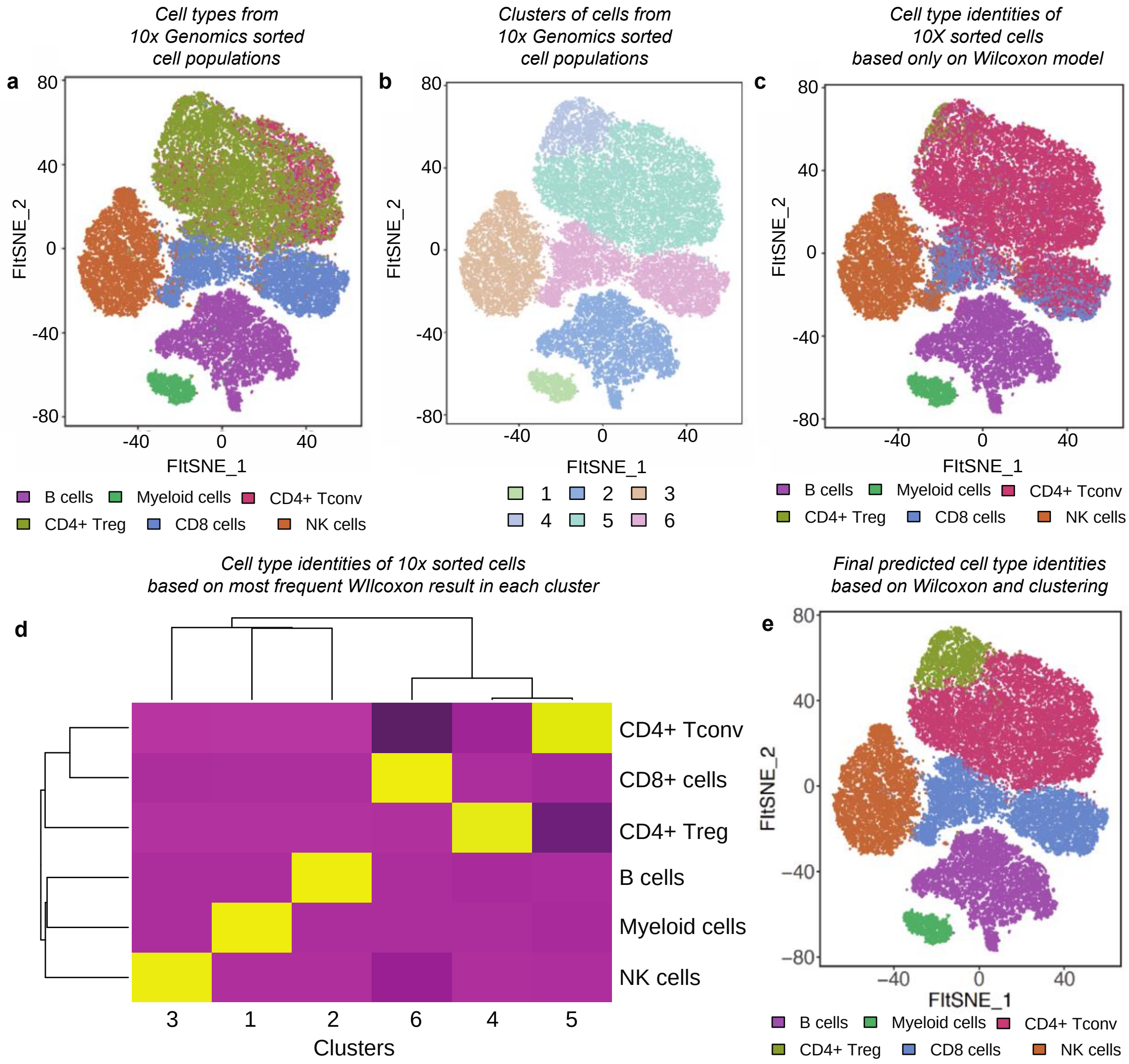


**f**

*Metrics of cell type predictions from Wilcoxon and clustering model using sorted cells from 10X Genomics*

| | B cells | Myeloid cells | CD4+ Tconv | CD4+ Treg | CD8+ cells | NK cells |
|---|---|---|---|---|---|---|
| Sensitivity | 1.000 | 0.987 | 0.976 | 0.928 | 0.990 | 0.985 |
| Specificity | 0.999 | 1.000 | 1.000 | 1.000 | 0.981 | 0.997 |
| Pos Pred Value | 0.997 | 0.996 | 1.000 | 1.000 | 0.926 | 0.987 |
| Neg Pred Value | 1.000 | 1.000 | 0.987 | 0.995 | 0.998 | 0.997 |
| Prevalence | 0.191 | 0.032 | 0.354 | 0.068 | 0.194 | 0.161 |
| Detection Rate | 0.191 | 0.031 | 0.346 | 0.063 | 0.192 | 0.158 |
| Detection Prevalence | 0.192 | 0.031 | 0.346 | 0.063 | 0.208 | 0.161 |
| Balanced Accuracy | 1.000 | 0.993 | 0.988 | 0.964 | 0.986 | 0.991 |

**Extended Figure 1: Validation of a combination Wilcoxon rank sum test and clustering based method for identification of cell types**. Publicly available data from sorted populations of immune cells was combined, and a classification algorithm was employed to identify cells types. **a.** FItSNE plot of combined purified B cells, CD14+ monocytes, CD4+ helper T cells, CD4+ Treg, CD8+ T cells, and CD4+ regulatory T cells. **b.** Same FItSNE plot as (**a**), but showing clustering results. Clusters were strongly associated with cell types of purified populations. CD4+ Treg, despite overlapping with CD4+ Tconv as a purified population, were strongly associated with cluster 5, suggesting that the sorted population of CD4+ Treg were mixed with CD4+ Tconv. **c.** Raw results from testing of the Wilcoxon rank sum from known cell populations. Individual cells were scored for enrichment of markers associated with each purified cell population. Some clusters were readily identifiable as pure populations using just the Wilcoxon based enrichment, but mixtures of T cells were evident. **d.** Heatmap showing the association between inferred lineage type from the Wilcoxon rank sum scores and the clusters. Each cluster largely consisted of a major lineage when looking at the aggregate Wilcoxon rank sum test across clusters. **e.** Inferred cell types based on the association between Wilcoxon scores and clusters from (**d**). Cell type inference agreed strongly with results of clustering from (**b**). **f.** Confusion matrix comparing the inferred cell types to the ground truth. The sensitivity, specificity, and accuracy were between 0.93 and 1.0 for all lineages.

Extended data Fig. 2

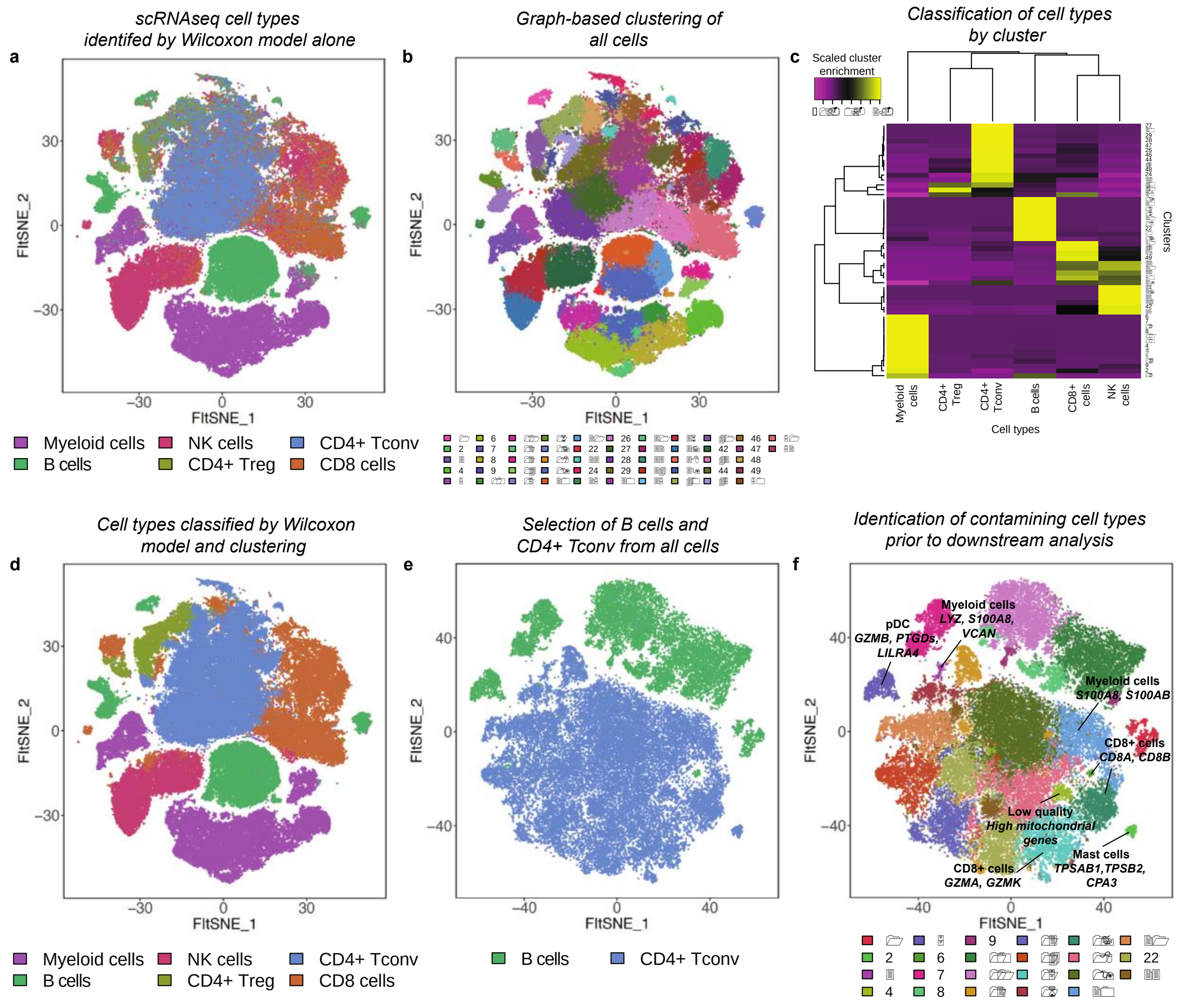

**Extended Figure 2: Identification of cell types from patients and controls using the combination Wilcoxon rank sum test and clustering based approach.**

**a**. Raw results from the Wilcoxon scores derived for each individual cell. Certain populations are highly accurately inferred from this first step, while others exhibit mixtures of cell types. **b.** Results of Louvian clustering revealed a total of 52 clusters from all the cells in the dataset. **c.** Heatmap showing the relationship between cluster and inferred cell types from (**a**) and (**b**). **d.** Results of the combination of Wilcoxon scoring and cluster association from (**a-c**). Major lineages are grouped together in FItSNE space, allowing the isolation of B cells and CD4+ Tconv for downstream analysis. **e.** B cells and CD4+ Tconv were bioinformatically isolated from D) and were projected in a new FItSNE space and colored by their inferred cell types. **f.** Identities of cell clusters were cross-checked by investigating differentially expressed genes across lineages. Contaminating lineages (i.e. those that are not B or CD4+ Tconv) and cell types that were not present in the training dataset (e.g. plasmacytoid dendritic cells [pDC], mast cells) were identified and removed, leaving only highly purified B and CD4+ Tconv for downstream analysis.

Extended data Fig. 3

**a**

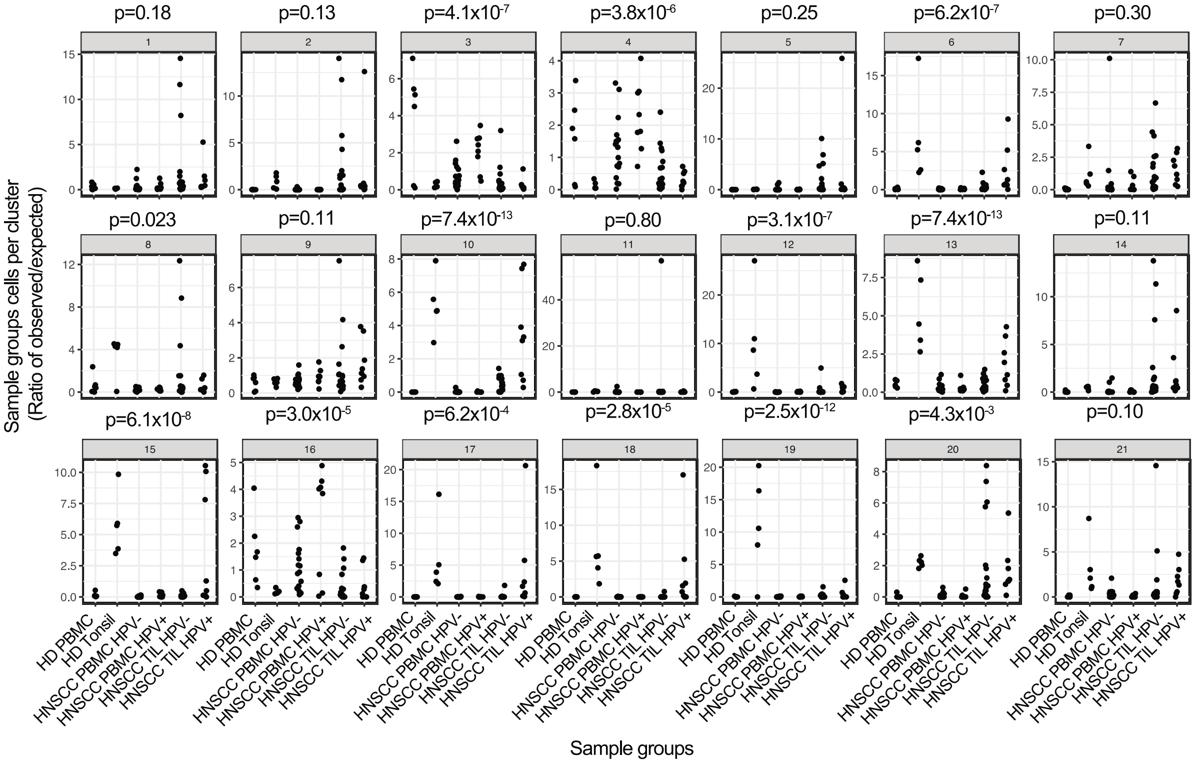


**b**

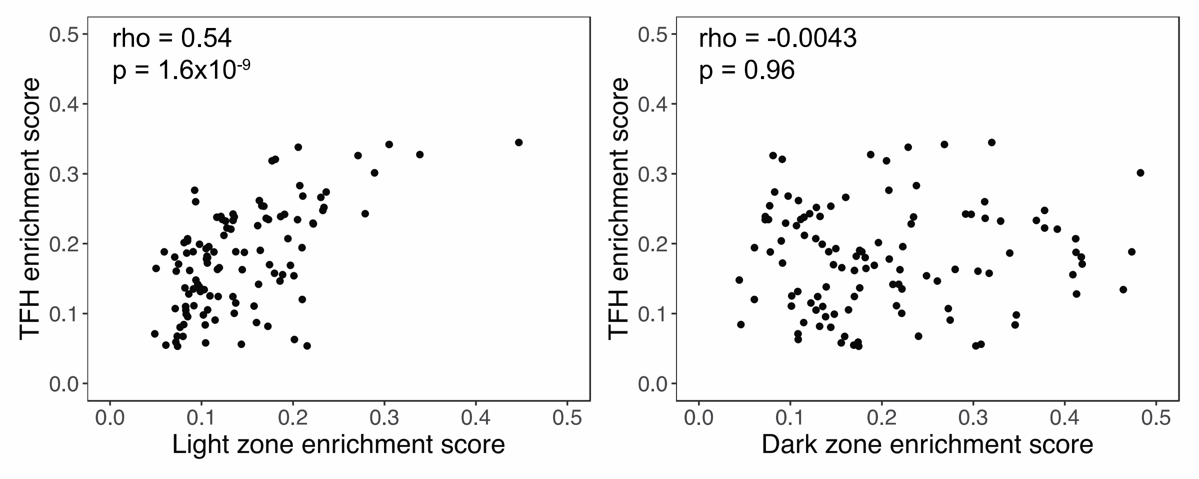

**Extended Figure 3: Statistical assessment of observed versus expected number of cells in each cluster by sample types.**

**a.** The expected frequency of cells in each cluster was calculated by the total cells in each cluster divided by the number of patients in the study. Cluster enrichment was calculated for each patient by dividing the observed over expected number of cells from each patient in each cluster. Analysis of variance was performed by calculating the cluster enrichment for each sample group in each cluster. Significant p values mean that one of more sample types deviate in a statistically significant way from the expected frequencies. **b.** Correlations between germinal center B cell enrichment scores and TFH enrichment scores derived from HNSCC patients in TCGA. Light zone B cells were significantly correlated with TFH cells (rho=0.54, p=1.6x10-9), but there was no relationship between dark zone B cells and TFH cells (rho=-0.0043, p=0.96).

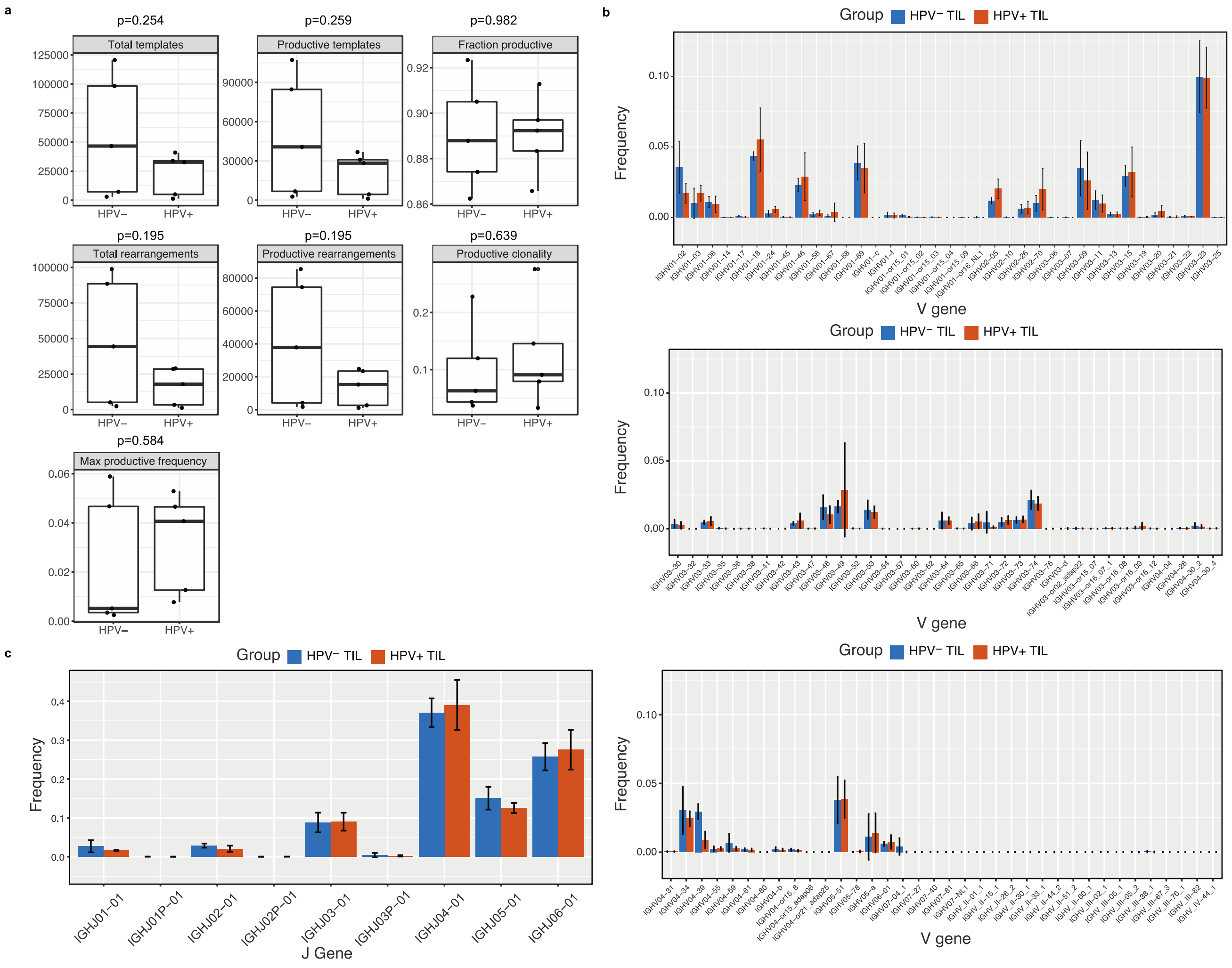
a
p=0.254
Total templates
p=0.259
Productive templates
p=0.982
Fraction productive
p=0.195
Total rearrangements
p=0.195
Productive rearrangements
p=0.639
Productive clonality
p=0.584
Max productive frequency
HPV−
HPV+
b
Group HPV− TIL HPV+ TIL
Frequency
V gene
c
Group HPV− TIL HPV+ TIL
Frequency
IGHJ01−01
IGHJ01P−01
IGHJ02−01
IGHJ02P−01
IGHJ03−01
IGHJ03P−01
IGHJ04−01
IGHJ05−01
IGHJ06−01
J Gene

**Extended Figure 4: Adaptive BCR sequencing reveals no difference in clonality or other metrics between HPV– and HPV+ TIL.**

**a.** There were no statistically significant differences in BCR templates between HPV– and HPV+ HNSCC TIL. **b.** V gene usage was largely similar between HPV+ and HPV– TIL. **c.** J gene usage was also similar between BCRs from HPV– and HPV+ TIL.

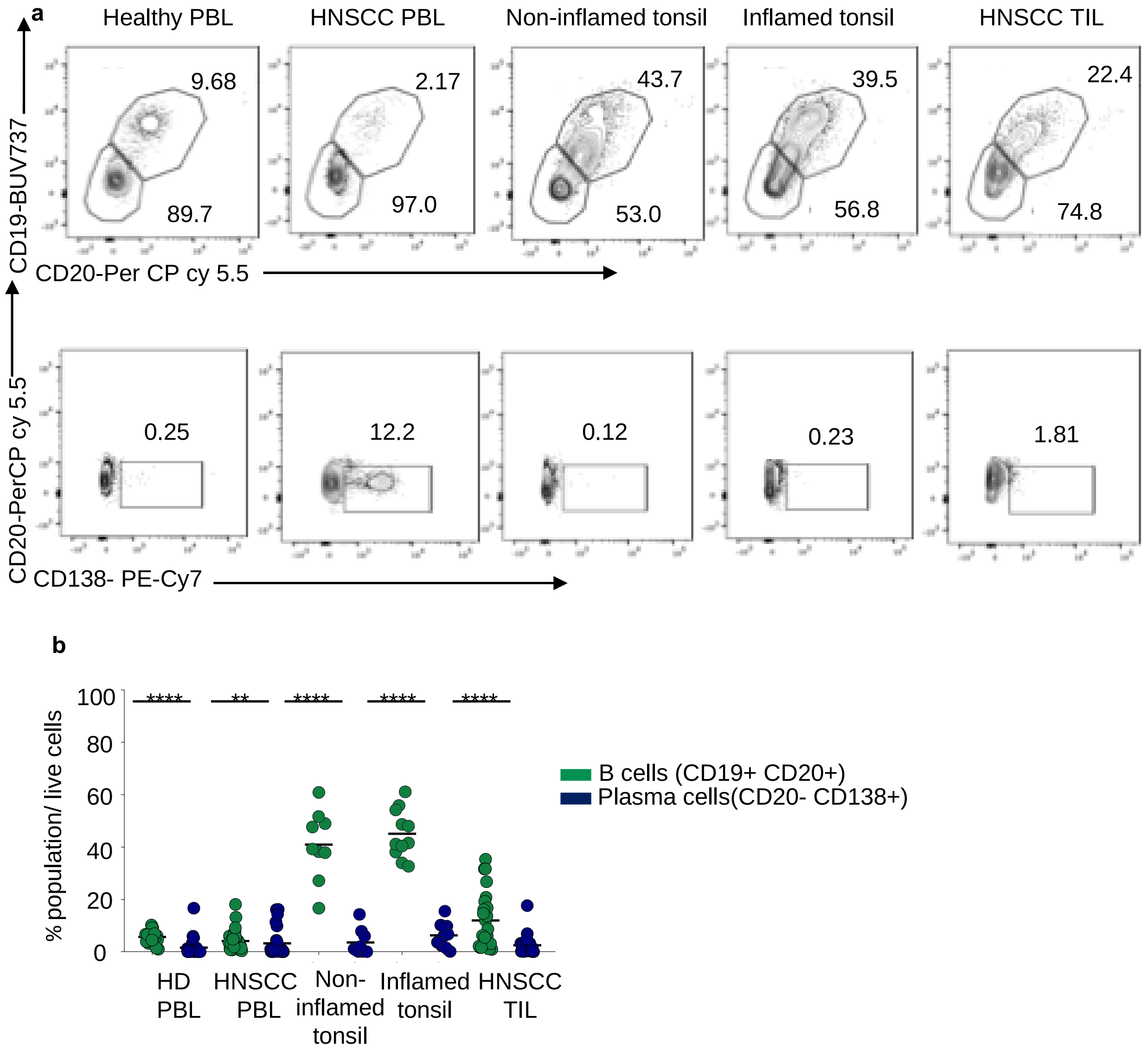

**Extended Figure 5: B cells are significantly increased compared to plasma cells in HNSCC patients.**

**a**. Representative flow plots for quantification of B cell frequency compared to plasma cell frequency from a separate cohort of tonsils, healthy PBMCs, HNSCC TIL, and HNSCC PBMC.

**b**. Scatter plot showing the frequency of B cells compared to plasma cells in healthy PBL (n=22), non-inflamed tonsils (n=9), inflamed tonsil (n=11), HNSCC tumor (n=23), HNSCC PBMCs (n=30). Statistical analysis by Students-T test (Mann-Whitney). ****P=<0.0001, **P=0.003

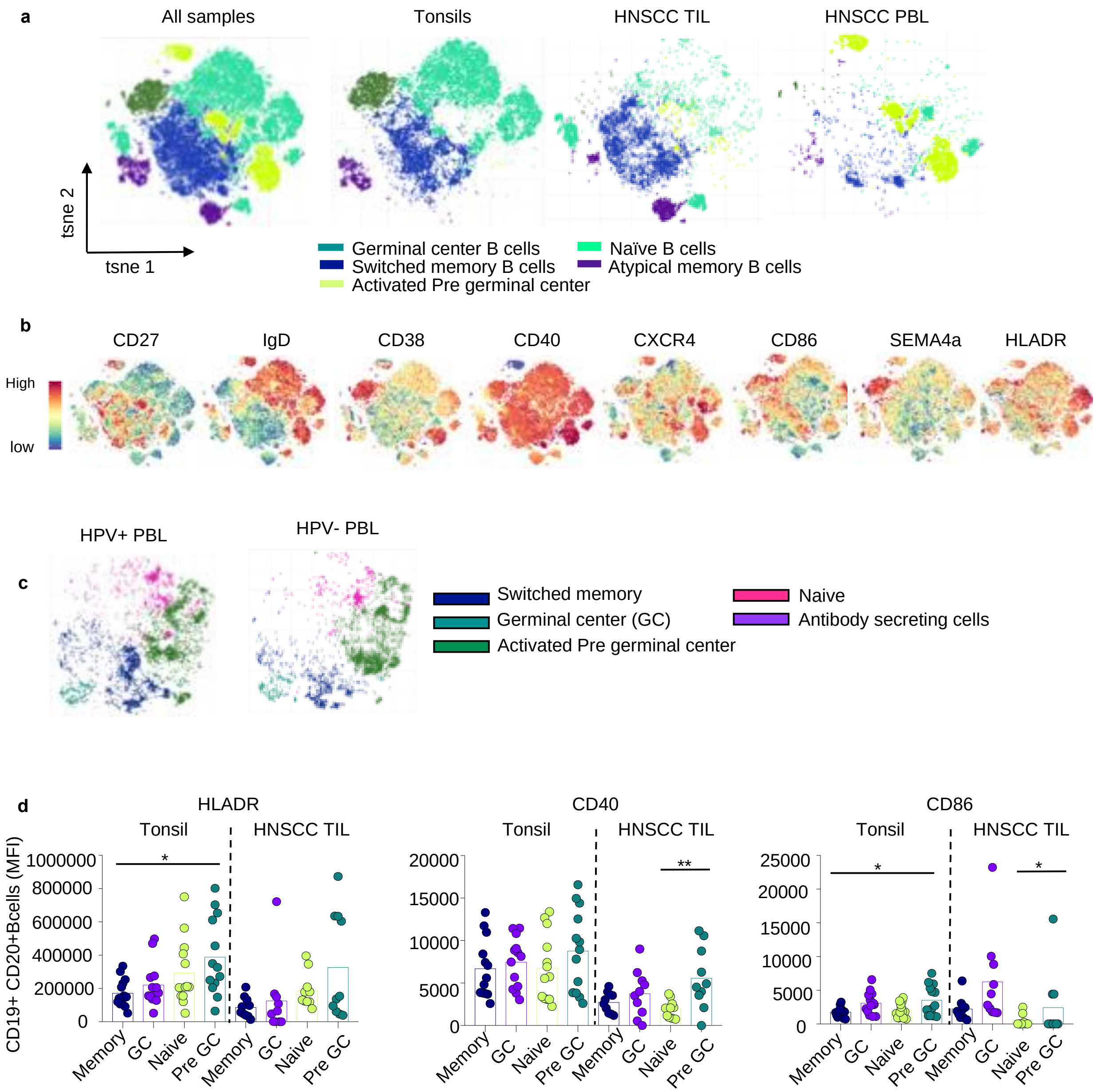
a
All samples
Tonsils
HNSCC TIL
HNSCC PBL
tsne 2
tsne 1
Germinal center B cells
Switched memory B cells
Activated Pre germinal center
Naïve B cells
Atypical memory B cells
b
CD27
IgD
CD38
CD40
CXCR4
CD86
SEMA4a
HLADR
High
low
c
HPV+ PBL
HPV- PBL
Switched memory
Germinal center (GC)
Activated Pre germinal center
Naive
Antibody secreting cells
d
HLADR
CD40
CD86
Tonsil
HNSCC TIL
CD19+ CD20+Bcells (MFI)
Memory
GC
Naive
Pre GC

**Extended Figure 6: Additional high dimensional analysis of HNSCC cohort 2.**

**a.** tSNE plots of all B cells collected from tonsils, HNSCC TIL and HNSCC PBMC analyzed using Cytofkit R program. Tonsils (n=1 sleep apnea patient, n=5 tonsillitis patients), HNSCC TIL (n=8), HNSCC PBMC (n=5). Cells are colored based on 5 populations identified using R phenograph. CD27, IgD, CD38, CXCR4, CD86, IgM surface markers were used to identify the 5 clusters**. b.** Individual feature plots demonstrating expression level of canonical markers used to identify B cell subpopulations. **c.** tsNE plots of B cell subsets in HNSCC PBL from HNSCC cohort used in figure 2a **d.** Bar plot showing mean fluorescent intensity of HLADR, CD86, CD40 on B cell subsets. Statistical analysis by one-way ANOVA followed by Tukeys multiple comparison. *P=0.03, **P=0.002.

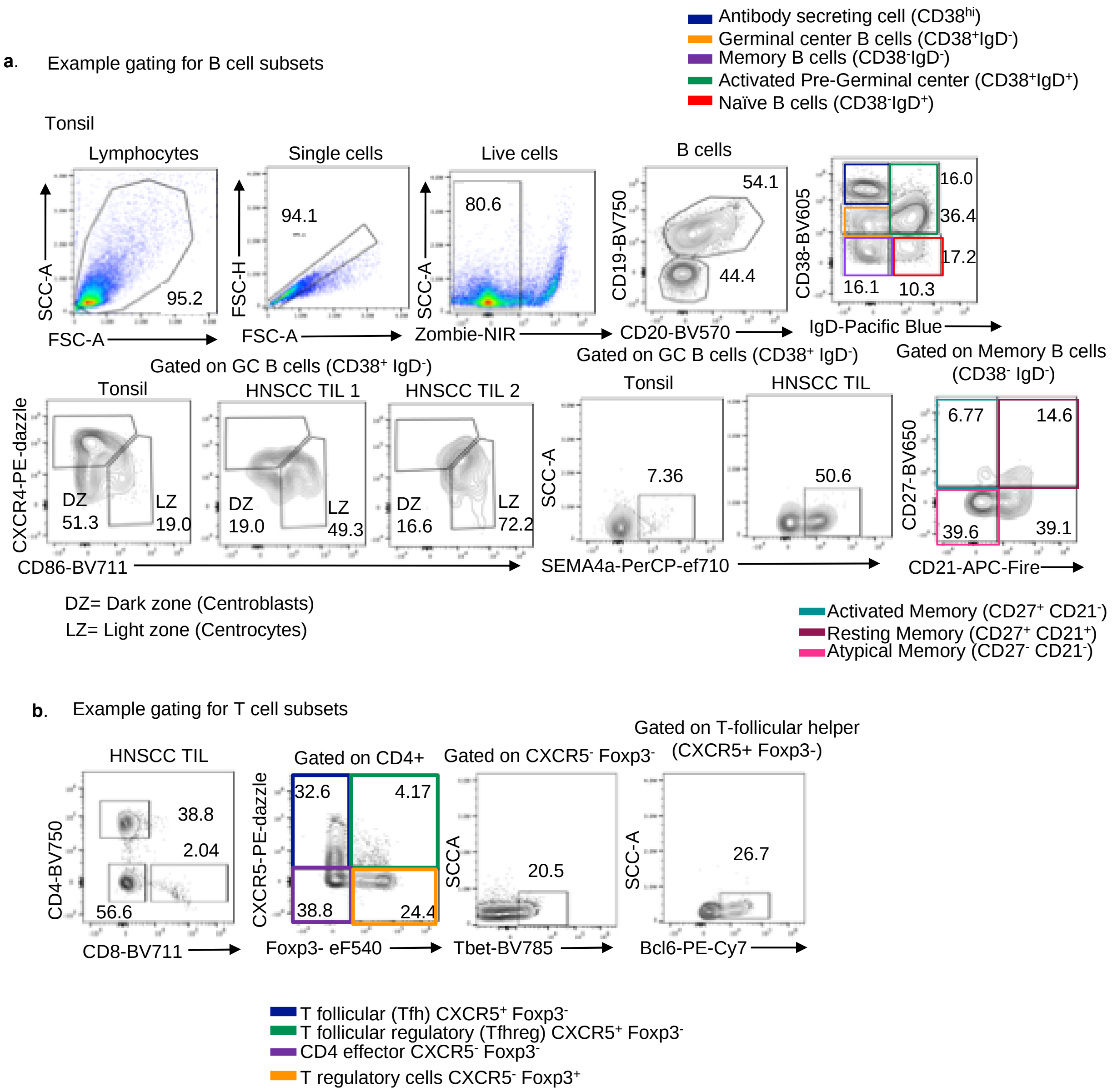
a. Example gating for B cell subsets
Antibody secreting cell (CD38hi)
Germinal center B cells (CD38+IgD-)
Memory B cells (CD38-IgD-)
Activated Pre-Germinal center (CD38+IgD+)
Naïve B cells (CD38-IgD+)
Tonsil
Lymphocytes
Single cells
Live cells
B cells
95.2
94.1
80.6
54.1
44.4
16.0
36.4
17.2
16.1
10.3
SCC-A
FSC-A
FSC-H
FSC-A
SCC-A
Zombie-NIR
CD19-BV750
CD20-BV570
CD38-BV605
IgD-Pacific Blue
Gated on GC B cells (CD38+ IgD-)
Tonsil
HNSCC TIL 1
HNSCC TIL 2
DZ 51.3
LZ 19.0
DZ 19.0
LZ 49.3
DZ 16.6
LZ 72.2
CXCR4-PE-dazzle
CD86-BV711
Gated on GC B cells (CD38+ IgD-)
Tonsil
HNSCC TIL
7.36
50.6
SCC-A
SEMA4a-PerCP-ef710
Gated on Memory B cells (CD38- IgD-)
6.77
14.6
39.6
39.1
CD27-BV650
CD21-APC-Fire
DZ= Dark zone (Centroblasts)
LZ= Light zone (Centrocytes)
Activated Memory (CD27+ CD21-)
Resting Memory (CD27+ CD21+)
Atypical Memory (CD27- CD21-)
b. Example gating for T cell subsets
HNSCC TIL
38.8
2.04
56.6
CD4-BV750
CD8-BV711
Gated on CD4+
32.6
4.17
38.8
24.4
CXCR5-PE-dazzle
Foxp3- eF540
Gated on CXCR5- Foxp3-
20.5
SCCA
Tbet-BV785
Gated on T-follicular helper (CXCR5+ Foxp3-)
26.7
SCC-A
Bcl6-PE-Cy7
T follicular (Tfh) CXCR5+ Foxp3-
T follicular regulatory (Tfhreg) CXCR5+ Foxp3-
CD4 effector CXCR5- Foxp3-
T regulatory cells CXCR5- Foxp3+

**Extended Figure 7: Flow cytometry gating strategy for B cell and T cell profiling.**

**a.** Representative flow cytometry plots for analysis of samples stained with 25 parameter Cytek Aurora panel (**Figures 2 and 4**). **b**. Representative flow cytometry plots for analysis of samples stained with T cell panel (**Figure 2c**).
